# Mutation rates and adaptive variation among the clinically dominant clusters of *Mycobacterium abscessus*

**DOI:** 10.1101/2022.04.27.489597

**Authors:** Nicoletta Commins, Mark R. Sullivan, Kerry McGowen, Evan Koch, Eric J. Rubin, Maha Farhat

## Abstract

*Mycobacterium abscessus* (*Mab*) is a multi-drug resistant pathogen increasingly responsible for severe pulmonary infections. Analysis of whole genome sequences (WGS) of *Mab* demonstrates dense genetic clustering of clinical isolates collected from disparate geographic locations. This has been interpreted as supporting patient-to-patient transmission, but epidemiological studies have contradicted this interpretation. Here we present evidence for a slowing of the *Mab* molecular clock rate coincident with the emergence of phylogenetic clusters. We find that clustered isolates are enriched in mutations affecting DNA repair machinery and have lower spontaneous mutation rates *in vitro*. We propose that *Mab* adaptation to the host environment through variation in DNA repair genes affects the organism’s mutation rate and that this manifests as phylogenetic clustering. These results inform our understanding of niche switching for facultative pathogens and challenge the model of transmission as the major mode of dissemination of clinically dominant *Mab* clusters.

## Introduction

*Mycobacterium abscessus* (*Mab*) is an emerging, multi-drug resistant pathogen that is increasingly responsible for opportunistic pulmonary disease, most commonly in patients with underlying structural lung conditions such as cystic fibrosis (CF)^1^. *Mab* is divided into three subspecies: *M. a. abscessus*, *M. a. massiliense*, and *M. a. bolletii,* the former two being the most common clinical subspecies^2^. Infections with *Mab* are associated with accelerated decline in lung function and can result in severe disseminated disease and/or loss of eligibility for lung transplantation. *Mab* infections are difficult to treat, requiring a prolonged antibiotic regimen with high rates of treatment failure^3^.

Until recently, *Mab* infections were thought to be acquired independently by each patient from environmental reservoirs in soil or water. Large scale analyses of whole genome sequencing (WGS) of Mab clinical isolates have revealed that a large proportion of clinical isolates fall into several phylogenetically characterized clusters, often termed dominant circulating clones^4^. Isolates within these clusters have high core genome similarity that is not explained by point source outbreaks as they were sampled from diverse geographic origins. These observations have been interpreted as supporting widespread recent transmission, either through direct contact or indirectly through fomites, but a number of epidemiological studies have refuted person-to-person transmission as a major mode of *Mab* dissemination^5–11^. Reconciling these conflicting lines of evidence is important as they dictate public health practice and allocation of resources to protect vulnerable populations from infection. However, it is still unclear what role transmission plays in the emergence of *Mab* clusters.

While widespread transmission is one proposed explanation for the phylogenetic structure observed in *Mab*, it is important to consider other factors in the species’ evolutionary history that may contribute. For example, transient niche switching between environmental sources and the human host may introduce bottlenecks resulting in low diversity and effective population size. Another explanation is a change in mutation rates, either through changes in bacterial generation time or through genetic variation. A third consideration is clinical sampling, where particular *Mab* lineages that cause more frequent, severe and/or persistent disease are more likely to be isolated and sequenced. Consistent with the latter hypothesis, previous work has shown that isolates within clusters are more likely to contain point mutations associated with antibiotic resistance and are associated with greater bacterial burden and lung inflammation in mice^4^.

Interpreting tree structures in the context of population history is aided by phylogenetic dating, which translates genetic distances into time scale. Previous studies^4,9,12,13^ have estimated molecular clock rates in *Mab* but these have been limited to subpopulations within clusters. Here we report on molecular clock rate estimation for the deep branches of the *M. a. abscessus* phylogeny and for within-host isolates belonging to clusters. We provide evidence for an evolutionary slow-down coincident with the expansion of clinically dominant clusters and propose that the emergence of dense phylogenetic clustering in *Mab* may result from genetically encoded changes to the mutation rate. These results indicate that transmission is not the only plausible explanation for the dense clustering observed in the *Mab* phylogeny, and that more research is needed to clarify the transmission dynamics of this emerging pathogen.

## Results

### Phylogenetic analysis confirms the presence of large, dominant clusters (DCs) spanning multiple geographies

We identified 1,629 *M. abscessus* isolates with publicly available whole genome sequencing (WGS) data and associated dates of collection (Table S1). We excluded isolates with likely contamination or low-quality assemblies (Methods). Among the 1,461 isolates passing quality filters, the majority (80.1%) were derived from *in vitro* culture of pulmonary samples (Table S2). Of the 1,181 pulmonary isolates, 98.3% (1,161/1,181) were collected from patients with cystic fibrosis (CF), while the rest were collected from patients with no documented diagnosis. The dataset represented 483 patients with unique patient identifiers, with 59.6% (288/483) of patients contributing a single isolate, and the remaining contributing between 2 to 24 isolates sampled over time. Isolates originated from nine countries and were collected between 1998-2017, except for the reference strain ATCC 19977 which was collected in 1957 (Figure 1B).

**Figure 1.**
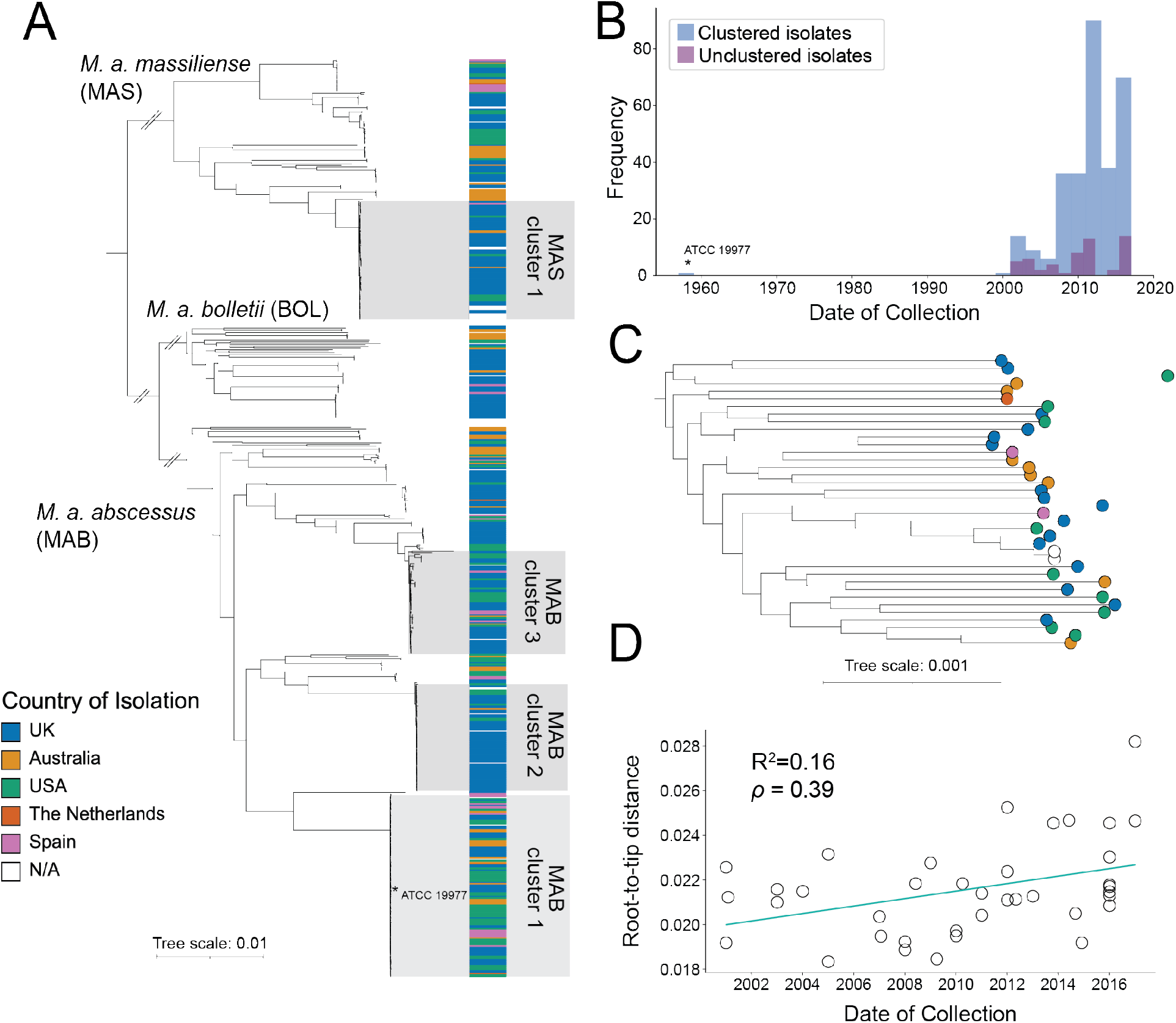
Dominant clusters of *M. abscessus* and the presence of temporal signal in internal branches. A) Species-wide phylogeny of *M. abscessus* representing one isolate per patient to avoid within-patient clustering of samples. Each subspecies tree was constructed independently using a subspecies specific reference genome. The four largest phylogenetic clusters are highlighted. B) Histogram showing the distribution of collection dates of all clustered isolates v. all unclustered isolates in *M. a. abscessus*. C) Pruned *M. a. abscessus* phylogeny representing 95% of the original subspecies tree diversity. C) Root-to-tip regression showing evidence of positive temporal signal in the pruned tree shown in (B).

The majority (91.7%) of samples had ≥98% average nucleotide identity (ANI) with one of three reference genomes (Table S3) representing each of the three Mab subspecies, consistent with the commonly used ANI threshold for bacterial subspecies. Isolates with <98% ANI with any subspecies were excluded from downstream analysis. The resulting dataset overrepresented clinically relevant subspecies of *M. abscessus*, with 93.9% belonging to either *M. a. abscessus* or *M. a. massiliense*. To better resolve true evolutionary relationships among samples, we defined and restricted analysis to the core genome for each subspecies, then further excluded predicted recombination events (Methods). We performed maximum likelihood (ML) phylogenetic analysis for each of the three *Mab* subspecies, using only the first isolate from each patient to focus on between-host diversity. The resulting subspecies trees confirmed the presence of multiple, globally distributed clades with dense phylogenetic clustering (Methods). We excluded clustering due to potential point source outbreaks by limiting to clusters containing isolates from more than one country and a minimum of three isolates for downstream analysis. In total, we confirmed 16 clusters (Figure S2A) including four clusters with > 50 isolates, which we refer to here as dominant clusters (DCs). Isolates within clusters comprised 76.4% of isolates included in the phylogenetic analysis. The median pairwise distance in *M. a. abscessus* DC 1 was 68 SNPs. For comparison, the mean pairwise distance across the subspecies trees were 5537, 4100, and 8034 SNPs for *M. a. abscessus*, *M. a. massiliense*, and *M. a. bolletii*, respectively. All four DCs contained isolates originating from three continents (North America, Europe, and Australia). Clustered isolates also spanned the full range of isolation times, including ATCC 19977, which was placed within *M. a. abscessus* DC 1 despite having been collected at least 40 years before any other isolates in the tree (Figure 1A–B).

### Coalescent analysis reveals a faster long-term evolutionary rate than observed in DCs

To aid in transmission inference and to reconstruct the evolutionary history of *Mab*, we performed a coalescent analysis using our SNP alignment. Previous work has reported molecular clock rates within *Mab* clusters, but to our knowledge no study has successfully dated any of the full *Mab* subspecies phylogenies. To evaluate the presence of temporal signal, we performed root-to-tip regression for each subspecies tree but in each case did not observe a positive correlation between sampling date and root-to-tip distance. We reasoned that the high degree of dense clustering in the *Mab* phylogeny may obscure the genetic-temporal signal. To test this hypothesis, we pruned each subspecies tree until it contained the minimal number of samples required to capture 95% of the genetic diversity in the original tree (Methods). Following pruning of the *M. a. abscessus* subspecies tree, root-to-tip regression demonstrated the presence of temporal signal (Figure 1C, D). Evidence of temporal signal in *M. a. abscessus* was further strengthened using a permutation test and a Bayesian evaluation of temporal signal (Figure S2C, Table S4).

We used a Bayesian implementation of coalescent analysis, BEAST^14^, to estimate the molecular clock rate of the pruned *M. a. abscessus* subspecies tree assuming a relaxed clock model, which allows each branch of the tree to have its own clock rate. To explore the effect of the tree prior on molecular clock estimation, we ran analyses using two different tree priors, assuming either a constant coalescent model, which assumes constant population size over time, or a Bayesian skyline coalescent model, which allows the population size to change over time. While the Bayesian skyline coalescent had a slightly better marginal likelihood, we found no significant difference between the clock rates estimated by the two models (Table 1, Figure 2C).

**Table 1.** Coalescent analysis results and model comparison

| Model | Marginal Log Likelihood (SD) | UCLD Rate (SNPs/site/year) | UCLD St. Dev. |
| --- | --- | --- | --- |
| Coalescent constant population | -7144301.64 (21.42) | $2.82 \times 10^{-6}$ [$1.33 \times 10^{-6}$, $4.83 \times 10^{-6}$] | 0.182 [0.135, 0.234] |
| Bayesian skyline coalescent | -7144118.76 (19.93) | $2.63 \times 10^{-6}$ [$1.16 \times 10^{-6}$, $4.0 \times 10^{-6}$] | 0.181 [0.138, 0.232] |

**Figure 2.**
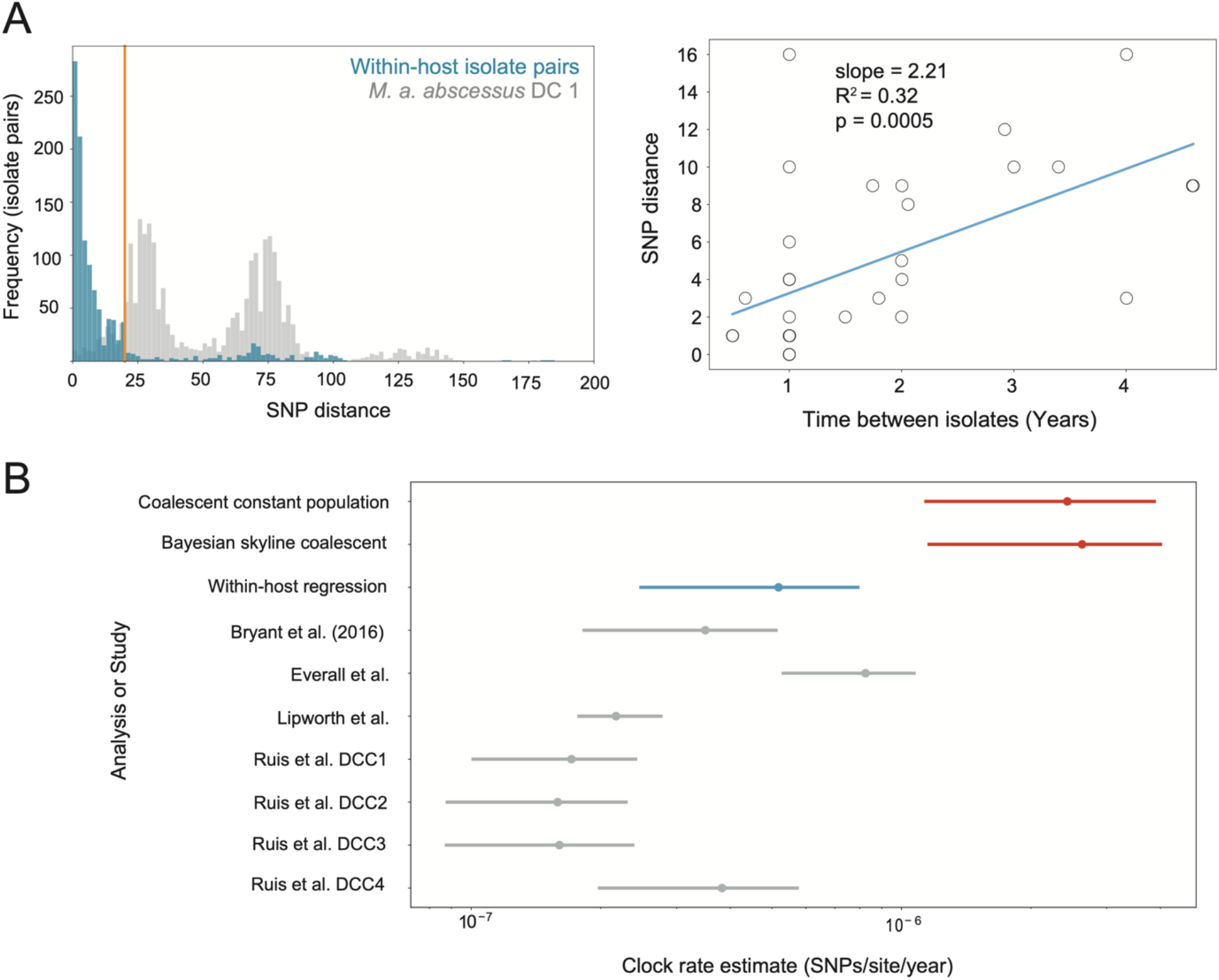
Longitudinally sampled within-host isolates and coalescent analysis support a slower clock rate within DCs. A) Clock rate estimation using isolate pairs sampled from the same patient over time. Left: distribution of SNP distances between all possible within-host isolate pairs compared to SNP distances between all possible isolate pairs within *M. a. abscessus* DC 1. The orange line represents the threshold of 20 SNPs used to exclude pairs representing independent infections. Right: regression of SNP distance on the time between isolates for each isolate pair. The slope of the regression line was 2.51 SNPs/year [1.24 – 3.78 95% CI], or roughly 5.93×10^−7^ SNPs/site/year [2.92 x10^−7^ – 8.9×10^−7^ 95% CI]. B) UCLD clock rate estimates (dots) for the three models used in this study (two coalescent models shown in red and one regression model shown in blue) compared to mutation rate estimates from the within-host regression (blue) and published clock rates (grey). Lines represent the 95% highest posterior density interval, except in the case of the within-host regression in which case the line represents the 95% confidence interval of the regression shown in (A).

Our estimates for the clock rate in the long, internal branches of the tree were approximately 10-fold faster than nearly all previously reported estimates of the clock rate within *Mab* clusters (Figure 2C). The clock rate estimate we obtain is sensitive to which samples are selected by Treemmer (Figure S3C). We therefore performed multiple runs using independent sets of isolates sampled by Treemmer and present the most conservative estimate we obtained from Treemmer sampling. Clock rate estimates were also robust to the algorithm choice or stringency of the recombination search procedure (Figure S3B). We next attempted to date the full (unpruned) *M. a. abscessus* phylogeny using the faster rate inferred from our coalescent analysis of the pruned tree. These chains suffered from poor convergence and sampled trees with a distorted topology, elongating the short branches and eliminating the densely clustered structure observed in the ML phylogeny (data not shown). We therefore hypothesized that the lack of temporal signal in the full *M. a. abscessus* dataset may result from a difference between the long-term clock rate and the more recent evolutionary rate within phylogenetic clusters.

### Analysis of longitudinally sampled within-host isolates supports a slower clock rate in DCs

To further investigate the differences in clock rate between long-term historical time scales and short-term contemporary time, we used pairs of isolates sampled from the same patient over time to estimate the mutation rate. We focused on estimating a within-host clock rate for *M. a. abscessus* to compare directly with the estimate from our coalescent analysis and because we had the most longitudinal samples for this subspecies. All isolate pairs belonged to one of the *M. a. abscessus* DCs. Because polyclonal *Mab* infection is common in CF patients, we first excluded any isolate pairs with a SNP distance > 20. This threshold was selected by comparing the distribution of all possible within-host isolate pairs to the distribution of pairwise SNP distances within *Mab* DCs (Figure 2A, Figure S4). Regressing the SNP distance between isolate pairs on the time between sample pairs yielded a mutation rate of 2.21 SNPs/year [1.06 – 3.37 95% CI], or 5.17×10^−7^ SNPs/site/year [2.48 x10^−7^ – 7.92×10^−7^ 95% CI]. This value is more similar to previously reported molecular clock rate estimates within *Mab* clusters^4,9,12,13^ than to those estimated in the long branches of the phylogeny (Figure 2C).

### Simulated ancestries support mutation rate slow-down over population size effects as the major driver of phylogenetic clustering

To test whether sampling from clades with low genetic diversity is sufficient to explain the degree of clustering observed in the ML phylogeny, or whether changes in the molecular clock rate are a better explanation for the observed phylognetic data, we performed coalescent simulations of ancestries with one clustered subpopulation (A) and one unclustered subpopulation (B). Under a range of effective population sizes (N_e_) and mutation rates (μ), we simulated ancestries through time with 1000 replicates per simulation. For each replicate we assessed the degree of phylogenetic clustering by measuring d_A_/d_B_ where d*_i_* is the mean pairwise SNP distance within subpopulation i. The total N_e_ (N_e_^T^), number of samples, and sampling dates for subpopulations A and B were drawn from the true values for *M. a. abscessus* DC 1 and from all “unclustered” isolates in our dataset, respectively (Figure 3A). We first tested whether a smaller population size for the clustered subpopulation A (N_e_^A^) could explain the degree of phylogenetic clustering observed. We estimated a range of realistic values for N_e_^A^ using BEAST by assuming a range of possible clock rates (Table S5). These rates spanned a published clock rate estimate for *M. a. abscessus* DC1 to a clock rate that is 50-fold higher, far exceeding the upper 95% highest posterior density (HPD) interval for the published value^15,16^. A schematic outlining this procedure is available as Figure S5. Under the the simulated conditions of equal mutation rates between subpopulation A and B, we only observed phylogenetic clustering to the extent observed in the clinical isolate phylogeny in the scenario where N_e_^T^ >> N_e_^A^ (Figure 3B, Figure S6). We next tested whether a change in the mutation rate at the root of supopulation A could better explain the observed phylogenetic structure in *Mab*. The simulated ancestries where N_e_^A^ and N_e_^B^ are fixed and the mutation rate slowdown is between 10- and 20-fold recapitulated the observed phylognetic structure (Figure 3C). This result was robust to changes in N_e_^A^ and N_e_^T^ across the 95% HPD for each parameter (Figure S7). These results indicate that for person-to-person transmission to drive phylogenetic clustering in *Mab*, the total N_e_ must be very large compared to the N_e_ of the clustered subpopulation, an unlikely scenario given the observed genetic data and sampling times. By contrast, a mutation rate change of 10- to 20-fold explains the degree of clustering observed in *Mab* under more realistic population sizes given these data.

**Figure 3.**
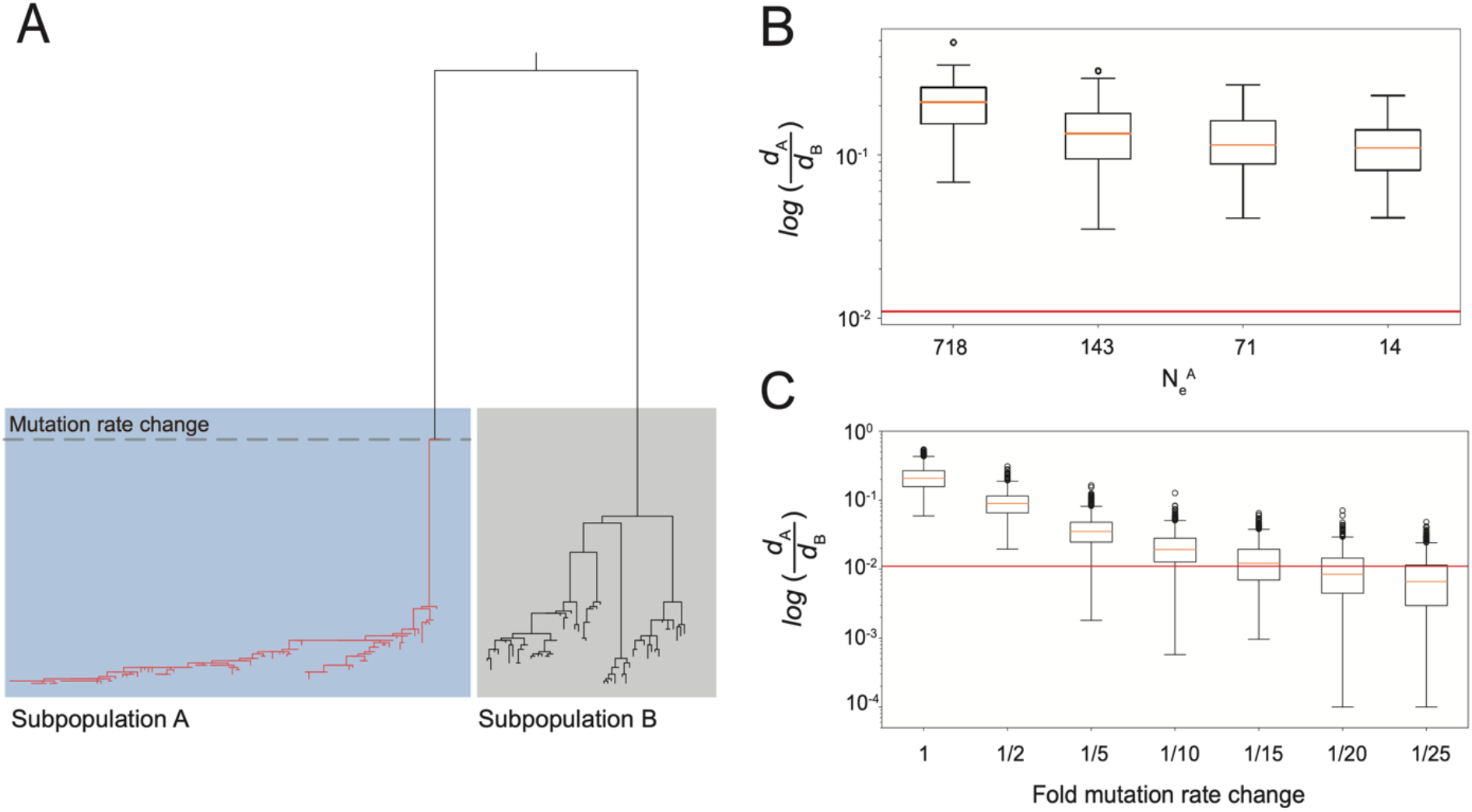
Simulated ancestries support a mutation rate slow-down coincident with DC emergence. A) Example neighbor joining (NJ) tree from the simulation shown in B. The number of samples and sampling dates for subpopulations A and B are drawn from real data from *M. a. abscessus* cluster 1 and from all unclustered samples, respectively. The dotted line crosses the common ancestor node for subpopulation A, and all branches descended from this node (shown here in red) are subject to the change in mutation rate. B) Boxplot showing the degree of phylogenetic clustering in subpopulation A relative to subpopulation B over a range of effective population sizes of subpopulation A (N_e_^A^). The simulation assumes that N_e_^T^=8000. C) Boxplot showing the degree of phylogenetic clustering in subpopulation A relative to subpopulation B over a range of mutation rate changes. The simulation assumes that N_e_^T^=8000 and N_e_^A^=718. In both B-C the degree of clustering is estimated with metric 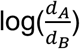 where d_i_ is the mean pairwise SNP distance among all sample pairs in subpopulation i. The red lines represent the clustering metric estimated using the observed ML phylogeny. Boxes extend from the lower to upper quartile values with a line at the median. Whiskers show the range of the data.

### UvrD/Rep family helicases are enriched in DCs in *M. a. abscessus*

To identify genetic variants under positive selection in *Mab* clusters, we searched for evidence of phylogenetic convergence in the core genome of *M. a. abscessus* with regions of recombination excluded. Briefly, we performed ancestral sequence reconstruction for the internal nodes of the subspecies tree and counted the number of independent arisals of each SNP (Methods). Of the 65,202 SNPs identified in *M. a. abscessus*, 34,813 (53.4%) of mutations occurred only once and were excluded. For the remaining SNPs, we ranked each by its enrichment in DCs relative to other regions of the *M. a. abscessus* tree (Figure 4A). The most significantly enriched variants in the *M. a. abscessus* DCs included homologs of several known or putative virulence factors as well as a number of cluster-enriched mutations in UvrD-like helicases *MAB_1054*, *MAB_3515c*, and *MAB_3516c* (Figure 4A, Table S6). While the function of these genes is not well characterized in *M. abscessus*, all three genes share homology with the UvrD/Rep family of helicases. This family of genes is involved in DNA repair and has been associated with growth and persistence *in vivo*^17^.

**Figure 4.**
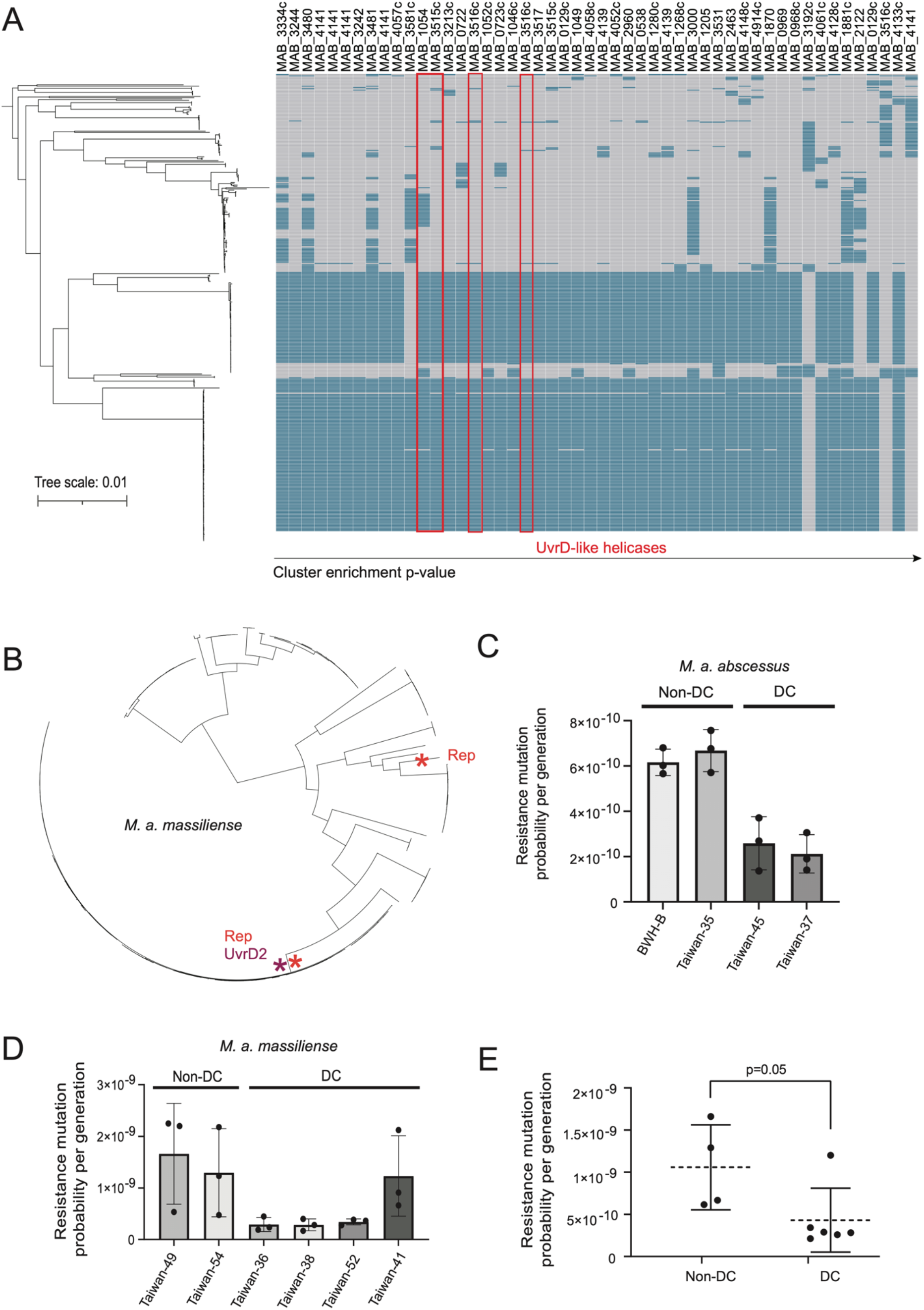
Dominant clusters are associated with a decrease in the spontaneous mutation rate *in vitro*. A) Subspecies phylogeny for *M. a. abscessus* (left) and corresponding heatmap representing the presence of the derived allele (blue) or ancestral allele (grey) for each nonsynonymous variant. Variants are ranked in ascending order by their p-value derived from Fisher’s tests for enrichment in the three largest clusters. The top 50 hits are shown. The genes where each variant is located are denoted. B) Subspecies phylogeny for *M. a. massiliense*. The branches where mutations have occurred in homologs of Rep and UvrD2 are shown with asterisks. Probability of acquiring an amikacin resistance mutation per generation in C) *M. a. abscessus* strains, D) *M. a. massiliense* strains, or D) in all strains tested that are part of a dominant cluster or not. Mean +/− SD is displayed. In C-D, n=3 biological replicates. In E, each point represents the average mutation probability for a single strain. p-value derived from two-tailed, unpaired t-test (t=2.258, df=8).

### Isolates in *Mab* DCs are associated with a lower spontaneous mutation rate

To test the association between phylogenetic clustering and spontaneous mutation rate, we selected *M. a. abscessus* and *M. a. massiliense* strains that were grouped within DCs or that were not present in those clusters. Each of the strains present in the DCs has the derived (minor) allele at four sites in the UvrD/Rep family genes *MAB_1054*, *MAB_3515c*, and *MAB_3516c*, and the strains outside of DCs possess the ancestral (major) allele at those sites (Table 2, Figure 4A). The strains used and their genotypes are described in Tables S7. The spontaneous mutation rate was estimated in all strains using a fluctuation assay^18^ adapted for use in *M. abscessus*. The fluctuation assay counts cells gaining resistance to the antibiotic amikacin during growth in culture without antibiotic. By normalizing the number of mutational events to the population size of the culture, the fluctuation assay estimates the spontaneous mutation rate per generation per the number of bases that can be mutated to confer resistance to amikacin. The strains chosen display similar growth rates (Figure S8A) and very low baseline resistance to amikacin (Table S8) which allows them to be compared using a fluctuation assay. Among *M. a. abscessus* strains, those present in DCs with the derived (minor) allele at the four UvrD/Rep loci had lower spontaneous mutation rates than those with the ancestral (major) allele (Figure 4C). A similar trend was observed in *M. a. massiliense* strains (Figure 4D). Further, the average mutation rate was significantly lower in isolates present in DCs across both subspecies (Figure 4E), consistent with the hypothesis that the emergence of dense clustering may be driven in part by changes in mutation rate.

**Table 2.**
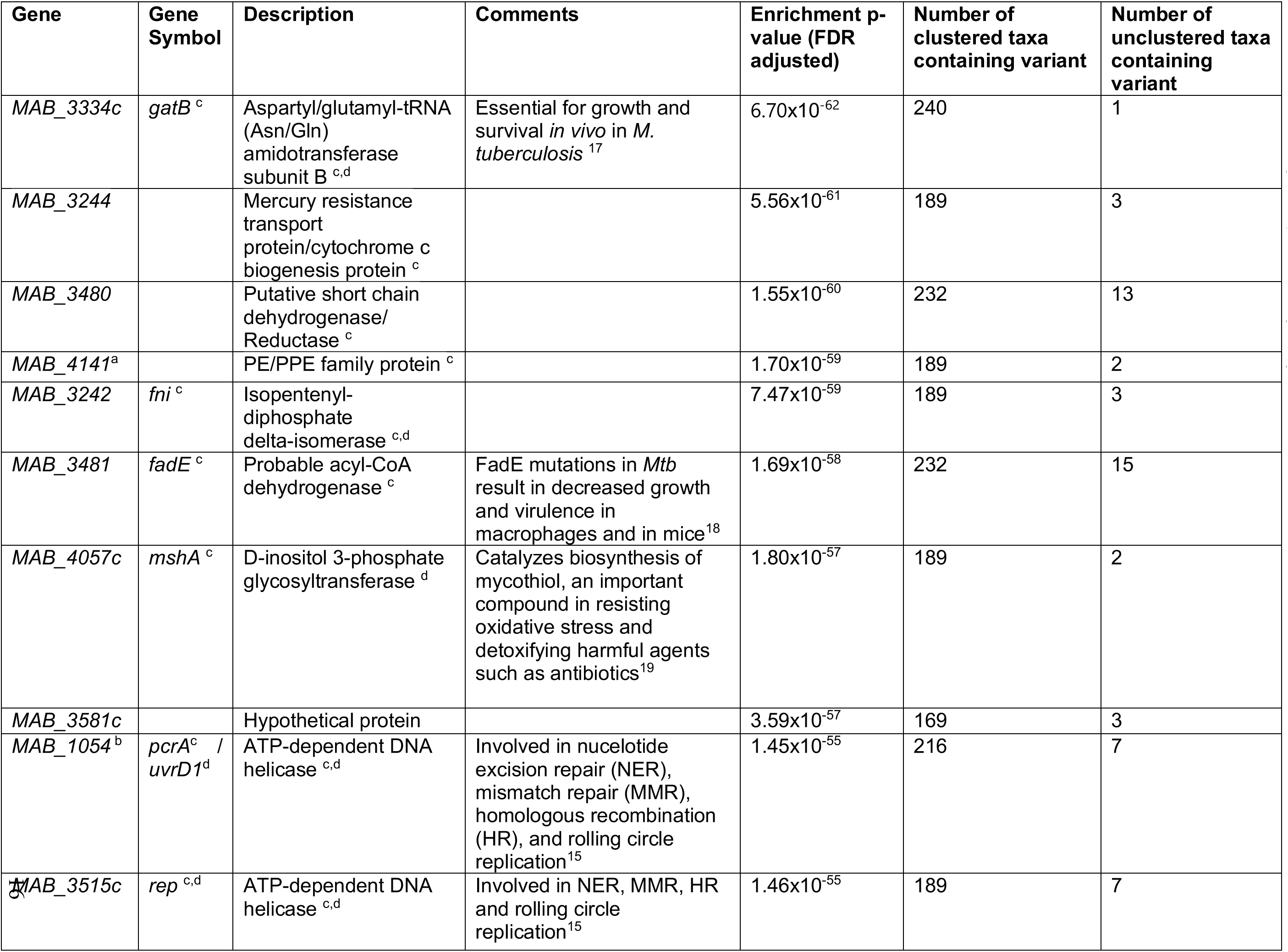
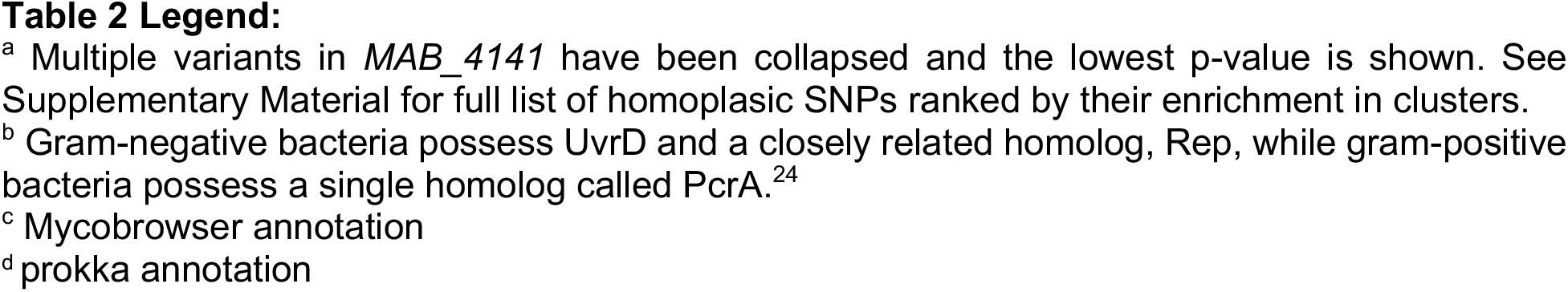
Summary of top 10 nonsynonymous variants enriched in DCs of *M. a. abscessus*

## Discussion

In this study we have pooled a large global sample of *Mab* genome sequences and examined evolutionary rates and positive selection with the goal of understanding the emergence of *Mab* clusters. We present evidence supporting a slower evolutionary rate within DCs compared to the long internal branches of the phylogeny and further data associating genetic variation in DNA repair genes with the slower evolutionary rate. We posit that changes to the mutation rate underlie the dense clustering in *Mab*, and that while other effects such as population or transmission bottlenecks and sampling bias may contribute to the low genetic diversity within clusters, they are not sufficient to explain the observed phylogenetic structure given the available data.

Previous studies have restricted molecular clock rate estimation to *Mab* clusters, most likely because of the lack of temporal signal at the subspecies-level. By pruning the *Mab* tree while preserving the majority of the subspecies diversity, we were able to measure a molecular clock rate for *Mab* spanning the time scale from the common ancestor of the subspecies to the present. Estimates of evolutionary rates are recognized to depend on the time scale of measurement. In such examples, the clock rate usually decreases with increasing time depth^18^. We observe the opposite effect where the clock rate increases with time depth. The pruned *Mab* tree most likely reflects an evolutionary process taking place predominantly in environmental reservoirs, in contrast to phylogenetic clusters where terminal branches reflect evolution within the human lung environment. We also note that our measurement of within-host mutation rate in *Mab* was independent of phylogenetic inference and the full temporal signal in our sample. Nevertheless, the measured rate within-host was comparable to the clock rate measured previously using coalescent analysis within clusters (Figure 2C).

The molecular clock rate is influenced by both the spontaneous mutation rate and the generation time of an organism. Changes in generation time could result from physiological constraints of the human lung environment, if for example, differences in nutrient availability change the growth rate in the lung compared to those living in environmental reservoirs. Generation times and spontaneous mutation rates can also be modified by genetic changes, for example to DNA replication or repair machinery. The observation that some lineages have acquired phenotypes that are beneficial in the host environment^4^ raises the possibility that genetic changes that underlie or associate with this adaptation affect mutation rate, either directly by affecting DNA replication or repair machinery, or indirectly by affecting the growth rate. Ruis et al. reported differences in mutational signatures along internal branches of the subspecies phylogeny compared to internal branches of DCs, supporting a change in the mutational process within some clades of the tree^19^.

We scanned the core genomes for variants enriched in the DCs in *M. a. abscessus*, the subspecies for which we have the most WGS data and for which there are the highest number of clusters. Notably, we observed several cluster-enriched mutations in UvrD-like helicases. UvrD is a superfamily 1 helicase that is involved in DNA repair and is widely conserved in gram-negative bacteria. UvrD1 in *M. tuberculosis* plays an important role both in nucleotide excision repair (NER) and in pathogenesis and persistence because its DNA repair activity confers tolerance to reactive oxygen intermediates (ROI) and reactive nitrogen intermediates (RNI)^20^. Mutations that promote the activity of UvrD1 and its homologs may therefore increase resistance to genotoxic stress in the context of the human lung environment, with the additional consequence of decreasing the spontaneous mutation rate. We also observed mutations in UvrD-like helicases occurring along the cluster-defining branch of the large *M. a. massiliense* cluster, further supporting the role of the NER pathway in *Mab* pathogenesis.

Our study has several limitations. Our dataset overrepresented clinical isolates from the UK where routine whole genome sequencing is part of *Mab* surveillance. Oversampling of some lineages and geographies may mean that we have not adequately captured the global *M. abscessus* diversity and that we may have focused on large phylogenetic clusters because of their geographic distribution and not because of their clinical significance. While our results support a faster molecular clock rate in the long internal branches of the phylogeny compared to clustered lineages, we are unable to distinguish between rate differences due to the spontaneous mutation rate from generation time effects. Differences in growth conditions across environments is difficult to quantify as *Mab* is ubiquitous in both natural and built environments, but nutrient availability likely differs greatly between natural reservoirs such as soil and water compared to man-made reservoirs such as showerheads. A healthy lung supports relatively little bacterial growth, but inflammation increases nutrient availability by promoting mucus production and vascular leakage into the airway^21^. Our analysis of variants associated with phylogenetic clustering is also limited in its ability to identify true causal genotypes or those that are enriched in clusters due to hitchhiking to other adaptive variants.

It is possible that our recombination removal procedure does not detect every mutation imported by recombination, and that unpurged recombination events may contribute to the high clock rate we measure in the pruned phylogeny. We believe any effect of recombination is likely minimal for several reasons: 1) recombination rates vary widely across branches, making unpurged recombination events more likely to erode temporal signal than to cause a systematic error in clock rate estimates (Figure S2A), and 2) our clock rate estimates were robust to choice of algorithm or to the stringency of recombination search (Figure S2B. Moreover, the algorithms we used to purge recombination events were also used in previous studies that published *M. abscessus* phylogenies.^4,9,12,13^ If these algorithms fail to purge enough recombination events to result in extreme distortion of clock rate estimates, they must also create distortion in phylogenies and the inference that follows from them. This strengthens the argument that phylogenetic clustering is not sufficient to infer widespread transmission.

While our study argues against transmission as the main driver of *Mab* dissemination and the emergence of DCs, these findings do not address the geographic distribution of Mab within clusters. It is possible that DC-associated variants such as the UvrD/Rep mutations described in this study are acquired ancestrally to the emergence of each DC, then circulated via contaminated household or medical equipment. This is similar to the model proposed by Bryant and colleagues, in which variation acquired along the branches ancestral to DCs creates large phenotypic changes, conferring a sudden increase in intracellular survival, tolerance to nitric oxide, and amikacin resistance.^23^ It is likely that emergence of clades with higher virulence potential involves acquisition of multiple genotypes including core genome mutations and horizontal gene transfer. Convergent evolution can also act in concert with the Bass Becking hypothesis that “everything is everywhere but the environment selects.” This could generate DCs that have a common core genome across many genetic loci and are prevalent in human infection across a wide range of geographies. Currently there is little publicly available sequencing data from environmental isolates. Isolating and sequencing *M. abscessus* from the environment could aid in understanding why clustered *Mab* isolates are detected in distant geographic locations. Future work could also compare genotypes and mutation rates in environmental samples compared to clinical isolates.

In summary, we present new evidence for changes in the molecular clock rate that contribute to the dense phylogenetic clustering observed in *Mab*. These results argue against person-to-person transmission as the primary driver of clustering in the *Mab* phylogeny. Continuing genomic surveillance is needed to understand how *Mab* may be adapting to human infection and to characterize the risk of transmission to at-risk patients. We argue further that the phylogeographic structure of *Mab* is influenced by its complex ecological history as an emerging facultative pathogen that should be considered carefully in future analysis of *Mab* genomic data. Future work should extend these analyses by examining differences in mutation rates and phenotypes between clinical isolates and environmental strains for evidence of host adaptation.

## Supporting information

Metadata table

All cluster enrichment data (all mutations)

Dominant cluster enrichment data (all mutations)

All cluster enrichment data (nonsynonymous mutations)

Dominant cluster enrichment data (nonsynonymous mutations)

## Acknowledgements

We thank members of the Farhat lab for feedback and discussion. We are grateful to Luca Freschi for advice on processing WGS data, calling variants, and phylogenetic inference; Maximillian Marin for input on calculating mappability pileup; and Roger Vargas for support in homoplasy counting. We also thank Wilder Wohns for discussions on implementing ancestry simulations with mutation rate changes through time. M.R.S. is a Merck Fellow of the Damon Runyon Cancer Research Foundation, DRG-2415-20. Computational resources and support were provided by the Orchestra High Performance Compute Cluster at Harvard Medical School, which is funded by the NIH (NCRR 1S10RR028832-01).

## Author Contributions

Conceptualization, N.C. and M.F.; Methodology – Computational analysis, N.C., E.K., and M.F.; Methodology – Experimental studies, M.R.S., K.M.; Investigation – Computational analysis, N.C.; Investigation – Experimental studies, M.R.S., K.M.; Data Curation, N.C.; Writing – Original Draft, N.C. and M.R.S.; Writing – Review & Editing, N.C., M.R.S., E.K., M.F.; Supervision, E.J.R., M.F.

## Methods

### Data Sources and Description

WGS data were curated from published studies^4,7,10,12,22–24^ and additional data were downloaded from the NCBI Sequence Read Archive (SRA). A search of “abscessus” was conducted on October 4, 2019 using the filters “DNA” and “Illumina”. Sequences were selected that had an associated date of collection with the year at minimum. Collection dates were obtained either from the literature, SRA metadata table, or provided by the authors. A summary of all samples used in this study is provided in Extended Data Table 1. Raw sequencing reads were downloaded using the NCBI SRA toolkit v2.9.6^25^.

### Quality control of Illumina reads

Reads were trimmed and filtered with trimmomatic v0.36^26^. To detect contaminated samples, the mean GC content of the reads were compared to a modelled normal distribution of GC content using fastqc v0.11.8^27^. If the sum of the deviations from the normal distribution represented more than 30% of the reads, the sequencing run was excluded from further analysis.

### Subspecies assignment and genome assembly

Multiple sequencing runs corresponding to one biological sample were combined into one file. Reads were then assembled *de novo* using SPAdes v3.11.1^28^. Assemblies were compared to a reference sequence for each of the three subspecies (Table S3) and a whole-genome based average nucleotide identity (gANI) score was calculated using fastANI v1.2^29^. The resulting alignments were assigned to a subspecies based on having gANI of at least 98% with exactly one reference strain. Isolates that did not have at least 98% ANI with any reference strain were excluded. No isolates matched more than one reference genome. After assigning isolates to a subspecies, reads were mapped to the corresponding reference genome using BWA MEM v0.7.17^30^. After alignment, isolates with <80% coverage of the reference genome at a depth of at least 20x or with missing data at >30% of sites were excluded.

### Variant calling and SNP filtering

Variants were called using Pilon v1.23^31^. We excluded ambiguous calls as wells as indels and MNVs. We also excluded calls with low coverage using a minimum depth of either 10% of the mean coverage, or 5, whichever was greater (default settings for Pilon –mindepth). We used bcftools v1.10.2^32^ to exclude calls with mapping quality or base quality scores < 20.

### Core genome inference

Unlike the professional mycobacterial pathogens *M. tuberculosis* or *M. leprae*, which have small accessory genomes, *M. abscessus* has a large, open accessory genome^33^. To define the core genome for phylogenetic analysis and to exclude loci that are highly divergent from the reference sequence, we masked all loci for downstream analysis where read depth was <20x in more than 5% of the isolates in our sample set. The remaining loci were compared to an estimate of the core genome calculated by Roary^34^, and the two methods yielded nearly identical results. We further excluded regions with average mapping quality scores or average based quality scores <20. Finally, we used GenMap v1.1^35^ to calculate mappability scores across each reference genome as defined in reference^36^. We then calculated a mappability pileup score for each base position as described in reference^37^ and excluded loci where the mappability pileup score was <95% from downstream analysis. In total, we excluded 14.4%, 13.5%, and 15.1% of the *M. a. abscessus*, *M. a. massiliense*, and *M. a. bolletii* reference genome length, respectively.

### Phylogenetic tree inference

A full-length sequence alignment was created using a custom script, masking loci excluded by our filtering criteria described above. Gubbins v2.4.1^38^ was used to predict and mask regions of variation produced by recombination from the full-length alignment. A recombination-free SNP alignment generated by Gubbins was then used to build a phylogeny for each subspecies using IQ-TREE v1.6.12^39^, using the ModelFinder Plus (-mfp) option and 1000 bootstrap replicates. Clusters were defined as previously described^4^ using a one-dimensional scanning statistic to identify clades with a distribution of branch lengths that is shorter than the distribution of branch lengths in the rest of the tree^41^.

### Coalescent analysis

Temporal signal was assessed using root-to-tip regression. In both *M. a. abscessus* and *M. a. massiliense* subspecies trees and all three major clusters, a slightly negative slope was observed, indicated lack of temporal signal. We recognized that the high degree of clustering and short time span over which the clustered isolates were sampled may confound the genetic temporal signal. To address this, we pruned the trees down to taxa that preserved 95% of the relative tree length (RTL) of the original trees using Treemmer v0.3^42^. A threshold of 95% eliminated most of the dense clustering, but preserved the long branches of the original tree (Figure 2B, Figure S2B). Using the pruned tree, we observed evidence of temporal signal in the MAB subspecies by regressing the root-to-tip distance for each sample against the date of collection. The linear regression was performed using the statsmodels package and the Pearson correlation coefficient was calculated using the pingouin package in Python. We generated a SNP alignment including only the taxa present in the pruned *M. a. abscessus* tree and used this as input for coalescent analysis using BEAST v2.6.3^42^. ModelTest-NG v0.1.6 ^43^ was used to select the site model GTR+4. MegaX^44^ was used to perform a maximum likelihood test to determine that a relaxed clock model is best suited for our data. This choice was confirmed using the nested sampling algorithm to compare marginal likelihoods for a strict v. relaxed clock model. We then used nested sampling to compare various tree priors. We used Bayesian Evaluation of Temporal Signal (BETS) to confirm the presence of temporal signal by comparing marginal likelihoods when the tip dates were used to those when all samples were assumed to be isochronous (Table S4). Based on marginal likelihood values, we ran both a coalescent constant population model and a Bayesian skyline coalescent model, assuming a relaxed clock and including the sampling dates for both models. We used the ML phylogeny of the pruned dataset as a starting tree. For each model we ran two independent chains for at least 100 million states to ensure convergence.

To test the effects of different recombination search procedures, we used Gubins v3.2.1 using 1) the default parameters and 2) more stringent search parameters by: decreasing the minimum number of SNPs to call a recombination to 2, increasing the significance threshold to p=0.10, and raising the trimming ratio to 1.5 to disfavor trimming of recombination edges. We also implemented recombination search using an alternative algorithm, ClonalFrameML^45^. We then used BEAST to estimate clock rates using each alignment, assuming a constant population tree prior (Figure S2B). To test the effects of which isolates are sampled by Treemmer, we ran Treemmer 10 times to obtain 10 independent samples. We then repeated our analysis using BEAST, again assuming a constant population tree prior (Figure S2C).

### Phylogenetic dating

To attempt to date the full MAB phylogeny, we ran both coalescent constant population and Bayesian skyline coalescent models assuming a relaxed clock without sampling dates. We set the mean UCLD clock rate to the values estimated from our coalescent analysis of the pruned tree. We ran each chain for at least 100 million states.

### Estimation of mutation rate within-host

We identified 90 patients with two or more *M. a. abscessus* isolates. Because coinfection with multiple *M. abscessus* clones is common in cystic fibrosis patients, we first sought to exclude any isolate pairs that represented distinct clonal infections. We selected a threshold of 20 SNPs by comparing the distribution of all possible within-host isolate pairs to the distribution of pairwise SNP distances within *Mab* clusters. We further excluded any isolates where >20% of the genome is predicted to be rearranged by recombination. For the remaining isolate pairs we regressed the pairwise SNP distance between the first and last isolates sampled from each patient on the time between the collection of each sample. The linear regression was performed using the Python statsmodels package v0.12.2. The slope of the regression line was used as an estimate of the number of SNPs accumulated over time.

### Ancestry simulations

We used msprime v1.0.2^46^ to simulate ancestries to model two subpopulations: (1) a “clustered” subpopulation (A) and (2) an “unclustered” subpopulation (B). We assumed a constant population size and initially set the total effective population size (N_e_) to 8000, the N_e_ estimated in our coalescent analysis of the pruned *Mab* tree. We chose a range of realistic N_e_ for subpopulation A, N_e_^A^ by running a separate coalescent analysis of *M. a. abscessus* cluster 1 using BEAST, assuming a constant population size, relaxed clock, and a range of UCLD clock rates spanning 1.7×10^−7^ SNPs/site/year, as previously reported for this cluster by Ruis et al. (2021) to 8.5×10^−6^ SNPs/site/year, a value well above the 95% HPD interval for our estimate of the clock rate in the pruned tree (Table S5). We assumed the N_e_ of subpopulation B to be: N_e_^B^ = N_e_^total^-N_e_^A^. The number of samples and sampling dates supplied to the model for subpopulations A and B are drawn from the true values for *Mab* cluster 1 and from all “unclustered” isolates in the tree, respectively. We assumed an initial mutation rate of 1×10^−9^ mutations/generation with a sequence length of 5Mb and a divergence time of 30,000 generations between subpopulations A and B. We also assumed the nucleotide subsitution model estimated using BEAST.

We first simulated a scenario where the mutation rate remains constant, varying N_e_^A^ over the range shown in Table S5. For each simulated ancestry we overlaid mutation events, supplying the model with a starting substitution rate of 1×10^−9^ mutations/generation, a nucleotide substitution model estimated using BEAST, and a sequence length of 5Mb. We then extracted the simulated genotypes and constructed a neighbor joining (NJ) tree using the Phylo module of the Python package biopython v1.79. The NJ tree was used to exclude any simulations where the two subpopulations did not exhibit reciprocal monophyly. We used the extracted genotypes to estimate the degree of clustering in each tree by comparing the mean pairwise SNP distance in subpopulation A (d_A_) to the mean pairwise SNP distance in subpopulation B (d_B_). For each simulated condition we performed 1000 replicates. To simulate ancestries with a mutation rate change, we repeated the above procedure conservatively assuming N_e_^A^ = 718. We modeled a per-generation mutation rate change starting at the common ancestor node for subpopulation A and ranging from 1-fold (no change) to 1/25-fold spanning a range of reasonable rate changes based on the range of clock rate estimates observed in this study and reported in the literature (Figure 2B). For all simulated conditions, we the simulation over a range of values for N_e_^total^ spanning the 95% HPD intervals for the N_e_ estimates using the Treemmer tree under both the constant coalescent and coalescent skyline tree priors.

### Scan for variants enriched in clinically successful clades

To search for variants associated with DCs, we first inferred the number of independent arisals in the MAB phylogeny for each SNP using SNPPar v1.0^46^. Briefly, SNPPar uses TreeTime to perform an ancestral sequence reconstruction at each node, then infers mutation events arising on each branch of the tree. We quantified the number of independent mutation events for each SNP. We considered only mutation events where the derived call was the minor allele and the ancestral call was the major allele. Because of the extreme phylogenetic clustering in *Mab*, we defined the major and minor alleles using the pruned alignment generated by Treemmer. For all homoplasic SNPs (occurring two or more times independently in the tree), we used Fisher’s exact test to quantify the enrichment of that variant within phylogenetic clusters and reported FDR-adjusted p-values to correct for multiple testing. We conducted two separate analyses: a) focusing on the three largest clusters in the *M. A. abscessus* tree (described in the main text), and b) considering all clusters identified by this approach. In each case we obtained similar results. For both analyses, full lists of variants ranked by their enrichment scores are provided in Supplementary Data. For each variant reported in Table 2, we manually examined SNPPar output to confirm that the variant was acquired independently along branches ancestral to at least 2 separate clusters. We also used SNPPar to identify all mutation events occurring on the branch ancestral to *M. a. massiliense* cluster 1.

### *M. abscessus* culture

*M. abscessus* clinical isolates used for fluctuation assays were isolated from patients in Taiwan and in Boston, MA, USA. All *M. abscessus* strains were cultured in Middlebrook 7H9 broth (271310, BD Diagnostics, Franklin Lakes, NJ, USA) with 0.2% (v/v) glycerol, 0.05% (v/v) Tween-80 (P1754, Sigma-Aldrich, St. Louis, MO, USA), and 10% (v/v) OADC (90000-614, VWR, Radnor, PA, USA). Proliferation rates were determined by inoculating 5 mL media with 500,000 colony forming units (cfu), then measuring OD600 at 24 and 48 hours post-inoculation. Proliferation rate was calculated as follows: proliferation rate in doublings per day = log_2_(OD600 at 48 hours / OD600 at 24 hours).

### Fluctuation Assay

Fluctuation assays were performed as described previously^17^ with adaptations for *M. abscessus*. *M. abscessus* strains were grown to saturation, then diluted to an OD600 < 0.01 and grown overnight until cultures reached an OD600 = 0.6-0.8. These cultures were used for fluctuation assays as well as determination of baseline amikacin sensitivity. Baseline amikacin sensitivity was tested by spreading dilutions of 10 million, 100 million, or 1 billion cfu on agar 7H10 (262710, BD Diagnostics) + OADC (90000-614, VWR) + 300 μg/mL amikacin sulfate (A2324, Sigma-Aldrich) plates. For fluctuation assays, 5000 cfu were plated in 1 mL media in each of 21 wells of a 24-well culture plate (10861-558, VWR), and initial cultures were plated for cfu on 7H10 + OADC to confirm number of cells in initial culture. 24-well plates were incubated shaking at 150 rpm for 6 days at 37°C. 18 wells of each strain were plated onto 6-well plates containing 5 mL 7H10 agar + OADC + 300 μg/mL amikacin sulfate in each well. Amikacin was used as the antibiotic selection for the assay because all tested strains had low baseline resistance (Table S8), which is required for the validity of the fluctuation assay. The remaining 3 wells from each strain were diluted and plated on 7H10 + OADC plates to enumerate average final cfu. Colonies on all agar plates were counted after 5 days incubation at 37°C. Mutation rate per generation per total number of bases that can be mutated to confer amikacin resistance was calculated using the Shinyflan application for the flan R package v0.9^17^ using the Maximum Likelihood estimation method, assumption of unknown fitness effects of mutations, the Exponential (LD model) distribution of mutant lifetime, and a Winsor parameter of 1024. One strain (BWH-E) was excluded from analysis due to highly variable results across replicates (Figure S6B). Mutation rates in clustered and unclustered samples were compared using a two-tailed, unpaired t-test with GraphPad Prism v9.3.1.

## Code Availability

Notebooks, snakemake piplines, custom scripts, conda environments, and BEAST XML files are available at: https://github.com/nicolettacommins/mab_mutation_rates_2022.

## Data Availability

Accession IDs for all publicly available data used in this study are available in the supplementary file: mabsc_all_sample_metadata.csv.

## References

1. Adjemian, J., Olivier, K. N. & Prevots, D. R. Epidemiology of Pulmonary Nontuberculous Mycobacterial Sputum Positivity in Patients with Cystic Fibrosis in the United States, 2010-2014. Ann. Am. Thorac. Soc. 15, 817–826 (2018).

2. Lee, M.-R. et al. Mycobacterium abscessus Complex Infections in Humans. Emerg. Infect. Dis. 21, 1638–1646 (2015).

3. Nessar, R., Cambau, E., Reyrat, J. M., Murray, A. & Gicquel, B. Mycobacterium abscessus: a new antibiotic nightmare. J. Antimicrob. Chemother. 67, 810–818 (2012).

4. Bryant, J. M. et al. Emergence and spread of a human-transmissible multidrug-resistant nontuberculous mycobacterium. Science 354, 751–757 (2016).

5. Bange, F. C., Brown, B. A., Smaczny, C., Wallace, R. J., Jr & Böttger, E. C. Lack of transmission of mycobacterium abscessus among patients with cystic fibrosis attending a single clinic. Clin. Infect. Dis. 32, 1648–1650 (2001).

6. Olivier, K. N. et al. Nontuberculous mycobacteria. I: multicenter prevalence study in cystic fibrosis. Am. J. Respir. Crit. Care Med. 167, 828–834 (2003).

7. Doyle, R. M. et al. Cross-transmission is not the source of new Mycobacterium abscessus infections in a multi-centre cohort of cystic fibrosis patients. Clin. Infect. Dis. (2019) doi:10.1093/cid/ciz526.

8. Sermet-Gaudelus, I. et al. Mycobacterium abscessus and children with cystic fibrosis. Emerg. Infect. Dis. 9, 1587–1591 (2003).

9. Lipworth, S. et al. Epidemiology of Mycobacterium abscessus in England: an observational study. The Lancet Microbe (2021) doi:10.1016/S2666-5247(21)00128-2.

10. Harris, K. A. et al. Whole-genome sequencing and epidemiological analysis do not provide evidence for cross-transmission of mycobacterium abscessus in a cohort of pediatric cystic fibrosis patients. Clin. Infect. Dis. 60, 1007–1016 (2015).

11. Tortoli, E. et al. Mycobacterium abscessus in patients with cystic fibrosis: low impact of inter-human transmission in Italy. Eur. Respir. J. 50, (2017).

12. Everall, I. et al. Genomic epidemiology of a national outbreak of post-surgical Mycobacterium abscessus wound infections in Brazil. Microb Genom 3, e000111 (2017).

13. Ruis, C. et al. Dissemination of Mycobacterium abscessus via global transmission networks. Nature Microbiology 1–10 (2021) doi:10.1038/s41564-021-00963-3.

14. Bouckaert, R. et al. BEAST 2.5: An advanced software platform for Bayesian evolutionary analysis. PLoS Comput. Biol. 15, e1006650 (2019).

15. Houghton, J. et al. Important role for Mycobacterium tuberculosis UvrD1 in pathogenesis and persistence apart from its function in nucleotide excision repair. J. Bacteriol. 194, 2916–2923 (2012).

16. Sinha, K. M., Unciuleac, M.-C., Glickman, M. S. & Shuman, S. AdnAB: a new DSB-resecting motor-nuclease from mycobacteria. Genes Dev. 23, 1423–1437 (2009).

17. Krašovec, R. et al. Measuring Microbial Mutation Rates with the Fluctuation Assay. J. Vis. Exp. (2019) doi:10.3791/60406.

18. Ho, S. Y. W., Shapiro, B., Phillips, M. J., Cooper, A. & Drummond, A. J. Evidence for time dependency of molecular rate estimates. Syst. Biol. 56, 515–522 (2007).

19. Huffnagle, G. B., Dickson, R. P. & Lukacs, N. W. The respiratory tract microbiome and lung inflammation: a two-way street. Mucosal Immunol. 10, 299–306 (2017).

20. Bryant, J. M. et al. Stepwise pathogenic evolution of Mycobacterium abscessus. Science 372, (2021).

21. Curti, E., Smerdon, S. J. & Davis, E. O. Characterization of the helicase activity and substrate specificity of Mycobacterium tuberculosis UvrD. J. Bacteriol. 189, 1542–1555 (2007).

22. Bryant, J. M. et al. Whole-genome sequencing to identify transmission of Mycobacterium abscessus between patients with cystic fibrosis: a retrospective cohort study. Lancet 381, 1551–1560 (2013).

23. Hasan, N. A. et al. Population Genomics of Nontuberculous Mycobacteria Recovered from United States Cystic Fibrosis Patients. bioRxiv 663559 (2019) doi:10.1101/663559.

24. Shaw, L. P. et al. Children with cystic fibrosis are infected with multiple subpopulations of Mycobacterium abscessus with different antimicrobial resistance profiles. Clin. Infect. Dis. (2019) doi:10.1093/cid/ciz069.

25. Download : Software : Sequence Read Archive : NCBI/NLM/NIH. https://trace.ncbi.nlm.nih.gov/Traces/sra/sra.cgi?view=software.

26. Bolger, A. M., Lohse, M. & Usadel, B. Trimmomatic: a flexible trimmer for Illumina sequence data. Bioinformatics 30, 2114–2120 (2014).

27. Babraham Bioinformatics - FastQC A Quality Control tool for High Throughput Sequence Data. https://www.bioinformatics.babraham.ac.uk/projects/fastqc/.

28. Prjibelski, A., Antipov, D., Meleshko, D., Lapidus, A. & Korobeynikov, A. Using SPAdes De Novo Assembler. Curr. Protoc. Bioinformatics 70, e102 (2020).

29. Jain, C., Rodriguez-R, L. M., Phillippy, A. M., Konstantinidis, K. T. & Aluru, S. High throughput ANI analysis of 90K prokaryotic genomes reveals clear species boundaries. Nat. Commun. 9, 5114 (2018).

30. Li, H. Aligning sequence reads, clone sequences and assembly contigs with BWA-MEM. arXiv [q-bio.GN] (2013).

31. Walker, B. J. et al. Pilon: an integrated tool for comprehensive microbial variant detection and genome assembly improvement. PLoS One 9, e112963 (2014).

32. Li, H. A statistical framework for SNP calling, mutation discovery, association mapping and population genetical parameter estimation from sequencing data. Bioinformatics 27, 2987–2993 (2011).

33. Choo, S. W. et al. Genomic reconnaissance of clinical isolates of emerging human pathogen Mycobacterium abscessus reveals high evolutionary potential. Sci. Rep. 4, 4061 (2014).

34. Page, A. J. et al. Roary: rapid large-scale prokaryote pan genome analysis. Bioinformatics 31, 3691–3693 (2015).

35. Pockrandt, C., Alzamel, M., Iliopoulos, C. S. & Reinert, K. GenMap: ultra-fast computation of genome mappability. Bioinformatics 36, 3687–3692 (2020).

36. Derrien, T. et al. Fast computation and applications of genome mappability. PLoS One 7, e30377 (2012).

37. Marin, M. et al. Benchmarking the empirical accuracy of short-read sequencing across the M. tuberculosis genome. Bioinformatics (2022) doi:10.1093/bioinformatics/btac023.

38. Croucher, N. J. et al. Rapid phylogenetic analysis of large samples of recombinant bacterial whole genome sequences using Gubbins. Nucleic Acids Res. 43, e15 (2015).

39. Minh, B. Q. et al. IQ-TREE 2: New Models and Efficient Methods for Phylogenetic Inference in the Genomic Era. Mol. Biol. Evol. 37, 1530–1534 (2020).

40. Harris, S. TreeGubbins. Preprint at https://github.com/simonrharris/tree_gubbins (2016).

41. Menardo, F. et al. Treemmer: a tool to reduce large phylogenetic datasets with minimal loss of diversity. BMC Bioinformatics 19, 164 (2018).

42. Darriba, D. et al. ModelTest-NG: A New and Scalable Tool for the Selection of DNA and Protein Evolutionary Models. Mol. Biol. Evol. 37, 291–294 (2020).

43. Kumar, S., Stecher, G., Li, M., Knyaz, C. & Tamura, K. MEGA X: Molecular Evolutionary Genetics Analysis across Computing Platforms. Mol. Biol. Evol. 35, 1547–1549 (2018).

44. Didelot, X. & Wilson, D. J. ClonalFrameML: efficient inference of recombination in whole bacterial genomes. PLoS Comput. Biol. 11, e1004041 (2015).

45. Kelleher, J. & Lohse, K. Coalescent Simulation with msprime. in Statistical Population Genomics (ed. Dutheil, J. Y.) 191–230 (Springer US, 2020). doi:10.1007/978-1-0716-0199-0_9.

46. Edwards, D. J., Duchêne, S., Pope, B. & Holt, K. E. SNPPar: identifying convergent evolution and other homoplasies from microbial whole-genome alignments. bioRxiv 2020.07.08.194480 (2020) doi:10.1101/2020.07.08.194480.

