## Supplemental Information for "Mutation rates and adaptive variation among the clinically dominant clusters of *Mycobacterium abscessus*"

### Supplementary Information

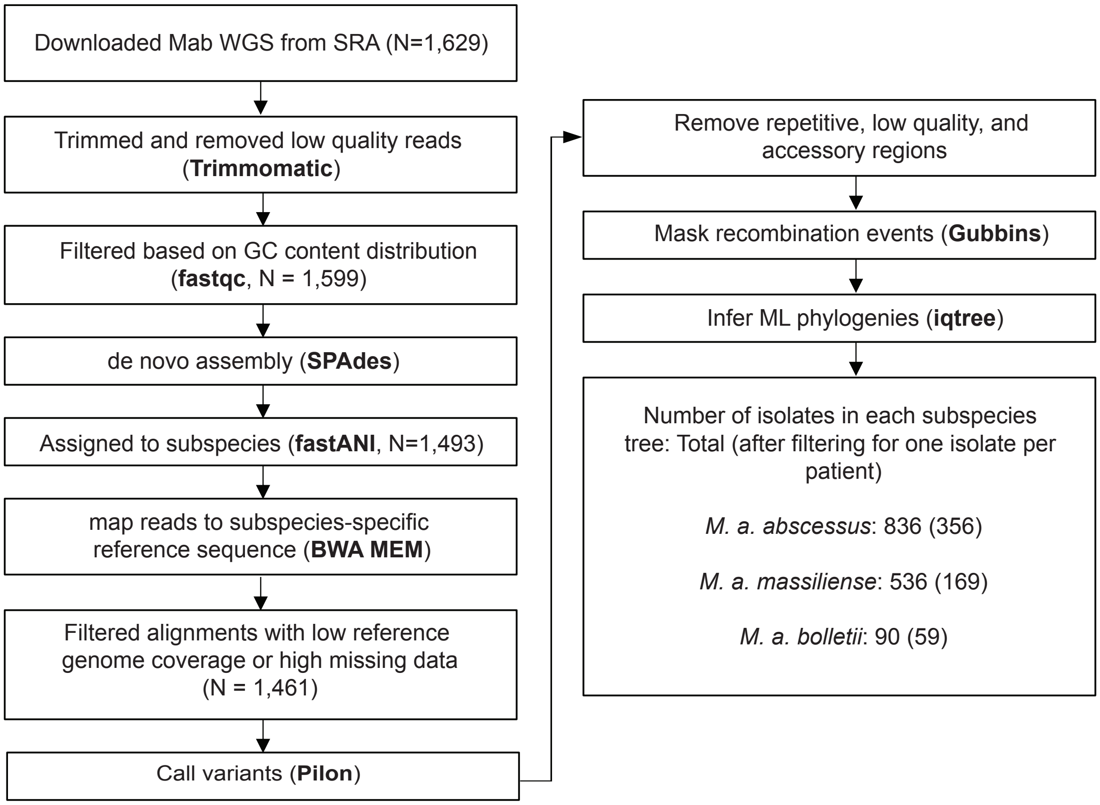


**Figure S1. Outline of methods for data acquisition, quality filtering, variant calling, and phylogenetic inference.** Software used are shown in bold. Other analysis steps were performed using custom scripts.

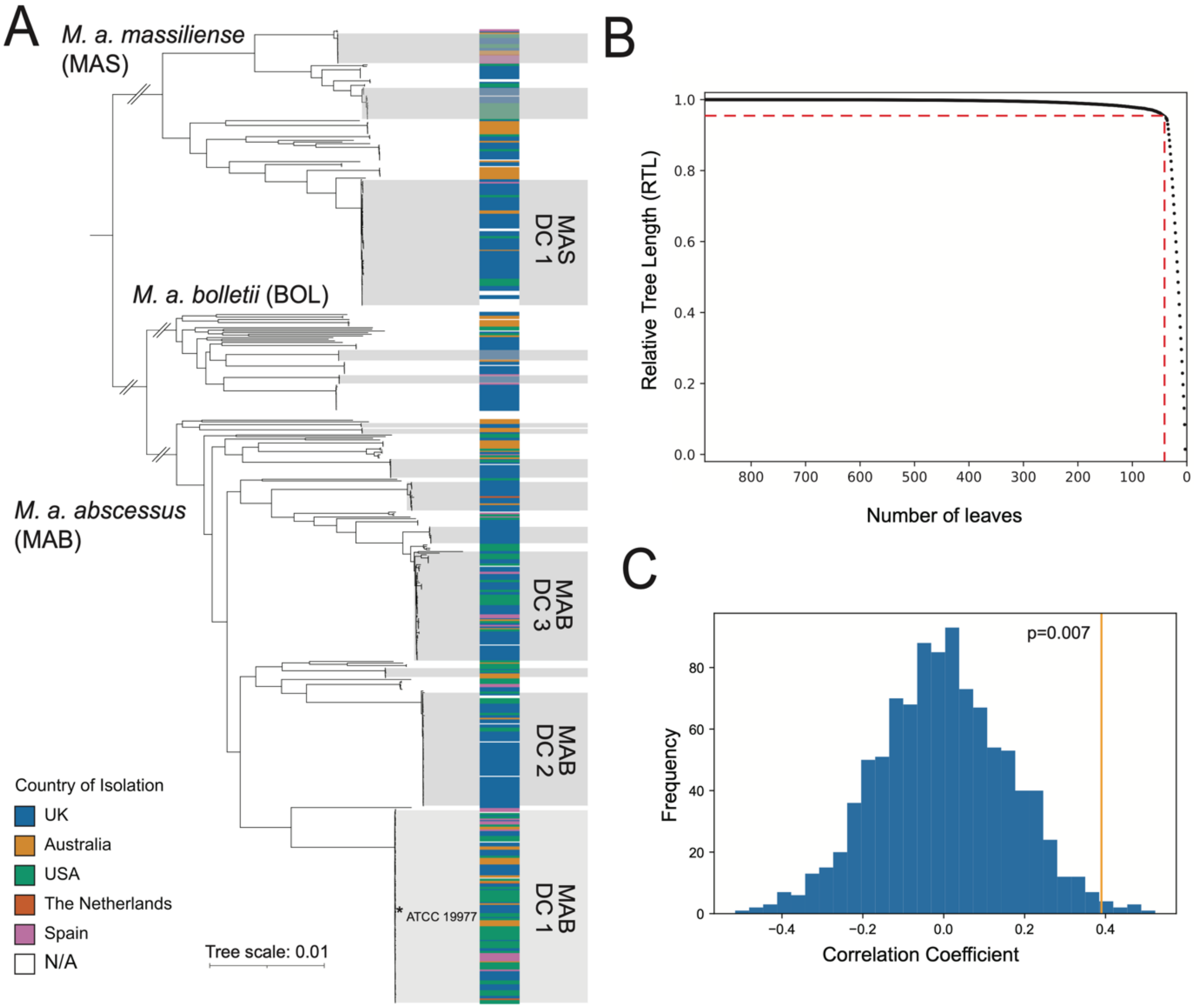


**Figure S2. *M. a. abscessus* clusters and inference of temporal signal (related to Figure 1).** A) Full species tree showing all clusters identified including DCs and non DC clusters. DCs are labeled. B) The relative tree length (RTL) v. the number of leaves remaining in the tree after each cycle of pruning with Treemmer. RTL is a measure of the diversity in the pruned tree relative to the original tree. The dotted red line indicates where the RTL is 95%, corresponding to 38 isolates. C) Permutation test for significance of temporal signal in the pruned *M. a. abscessus* tree. We randomly shuffled the dates of collection and root-to-tip distances from the pruned subspecies tree 1000 times. For each permutation we calculated Pearson's correlation between the collection dates and root-to-tip distances to estimate an empirical p-value = 0.007.

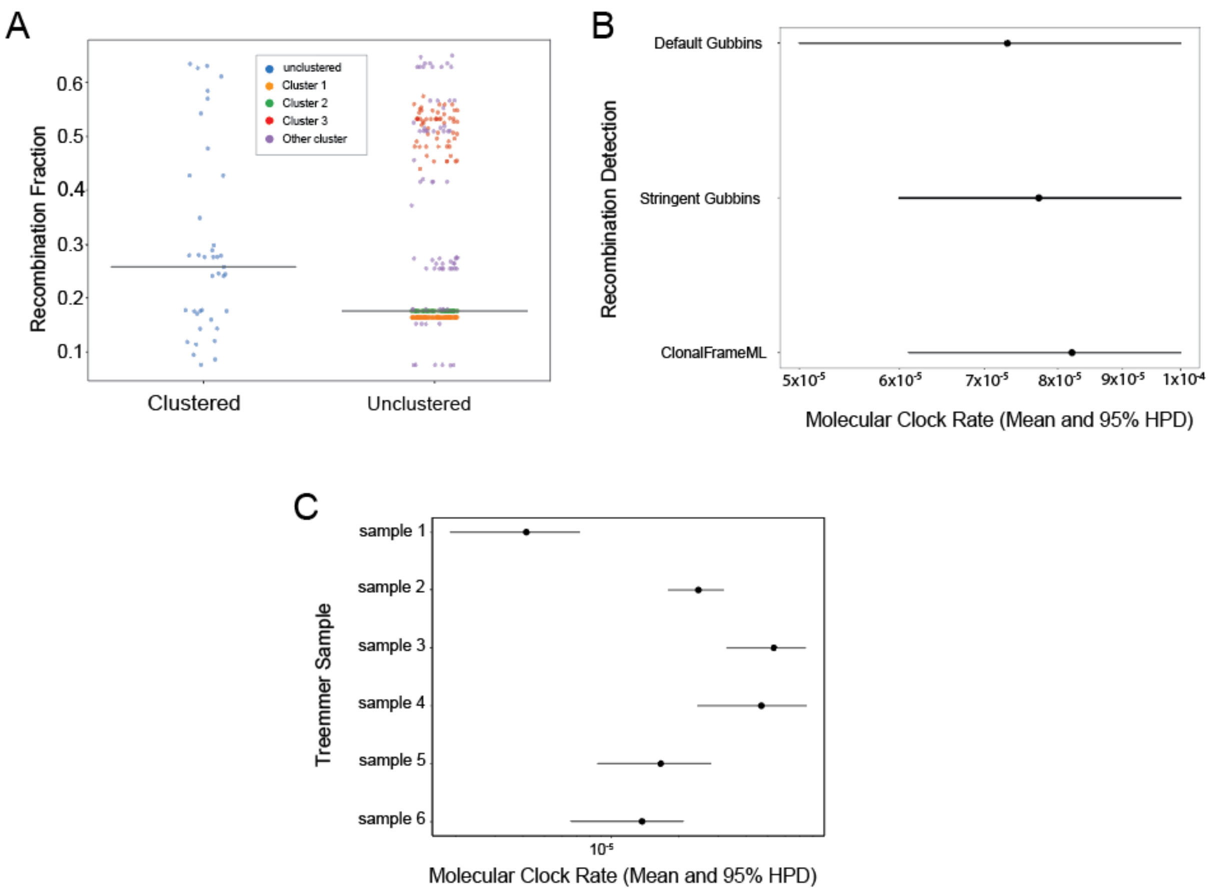


**Figure S3. Effect of recombination removal and sampling on clock rate estimates (Related to Figure 2).** A) Fraction of variation in each sample attributed to recombination by Gubbins. Horizontal lines indicate median values. B) Clock rate estimates after using the default Gubbins search (Methods) and ClonalFrameML. C) Clock rate estimates from independent Treemmer samples.

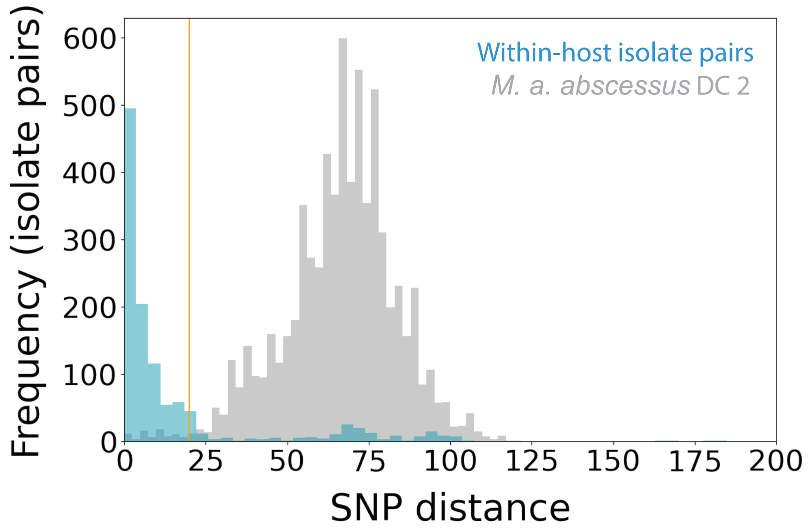


**Figure S4. SNP distance threshold for within-host isolate pairs (related to Figure 2A).** Distribution of SNP distances between all possible within-host isolate pairs in *M. a. abscessus* compared to SNP distances between all possible isolate pairs within *M. a. abscessus* DC 2.

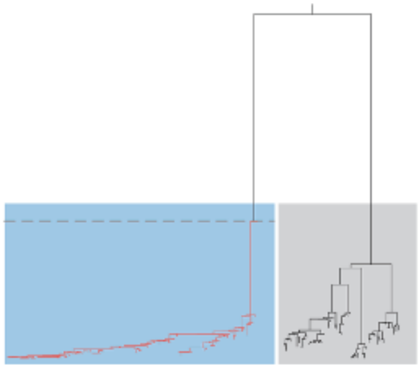

Subpop. A ("Clustered") Subpop. B ("Unclustered")

Determine a realistic range for ($N_e^A$):

| **Clock rate (SNPs/site/year)** | **Estimated $N_e^A$ [95% HPD]** |
|---|---|
| $1.7x10^{-7}$ * | 718.99 [575.06, 875.16] |
| $8.5x10^{-7}$ | 142.79 [116.38, 175.52] |
| $1.7x10^{-6}$ | 71.75 [57.57, 87.07] |
| $8.5x10^{-6}$ | 14.27 [11.47, 17.35] |

* Mutation rate estimated for MAB cluster 1 (Ruis et al, 2021)

Estimate effective population size ($N_e^B$) directly using alignment and sampling dates

Simulate ancestries (msprime) over range of values for $N_e^A$ and spanning the 95% HPD interval for $N_e^B$

**Figure S5. Schematic outlining procedure for selecting parameter values for ancestry simulations (Related to Figure 3).**

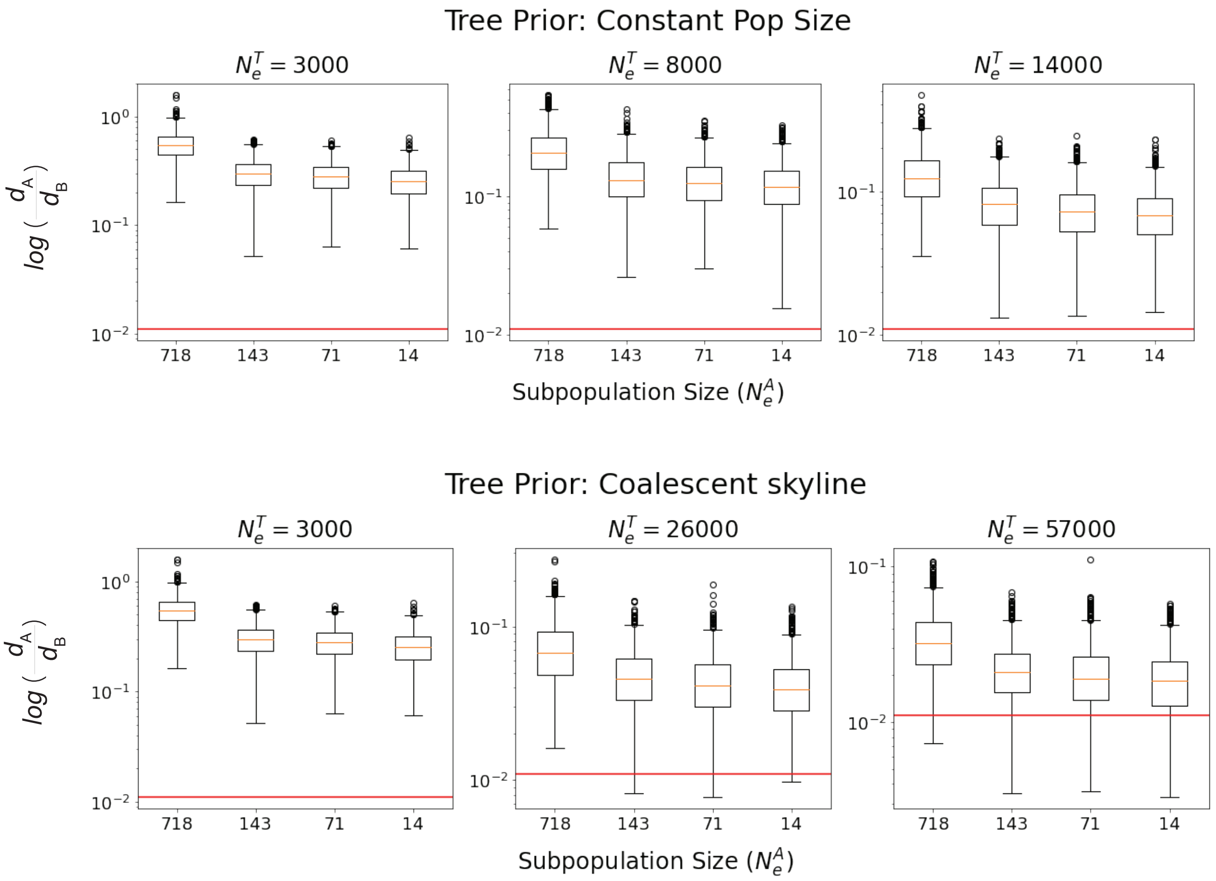


**Figure S6. Ancestry simulations across a range of possible population sizes (related to Figure 3A).** Boxplots showing the degree of phylogenetic clustering in subpopulation A relative to subpopulation B (Figure 3A) over a range of effective population sizes of subpopulation A ($N_e^A$) and a range of total effective population sizes ($N_e^T$) spanning the 95% confidence intervals estimated using BEAST under a constant population size model (top) and a Bayesian coalescent skyline model (bottom). The top center panel recapitulates figure 3B.

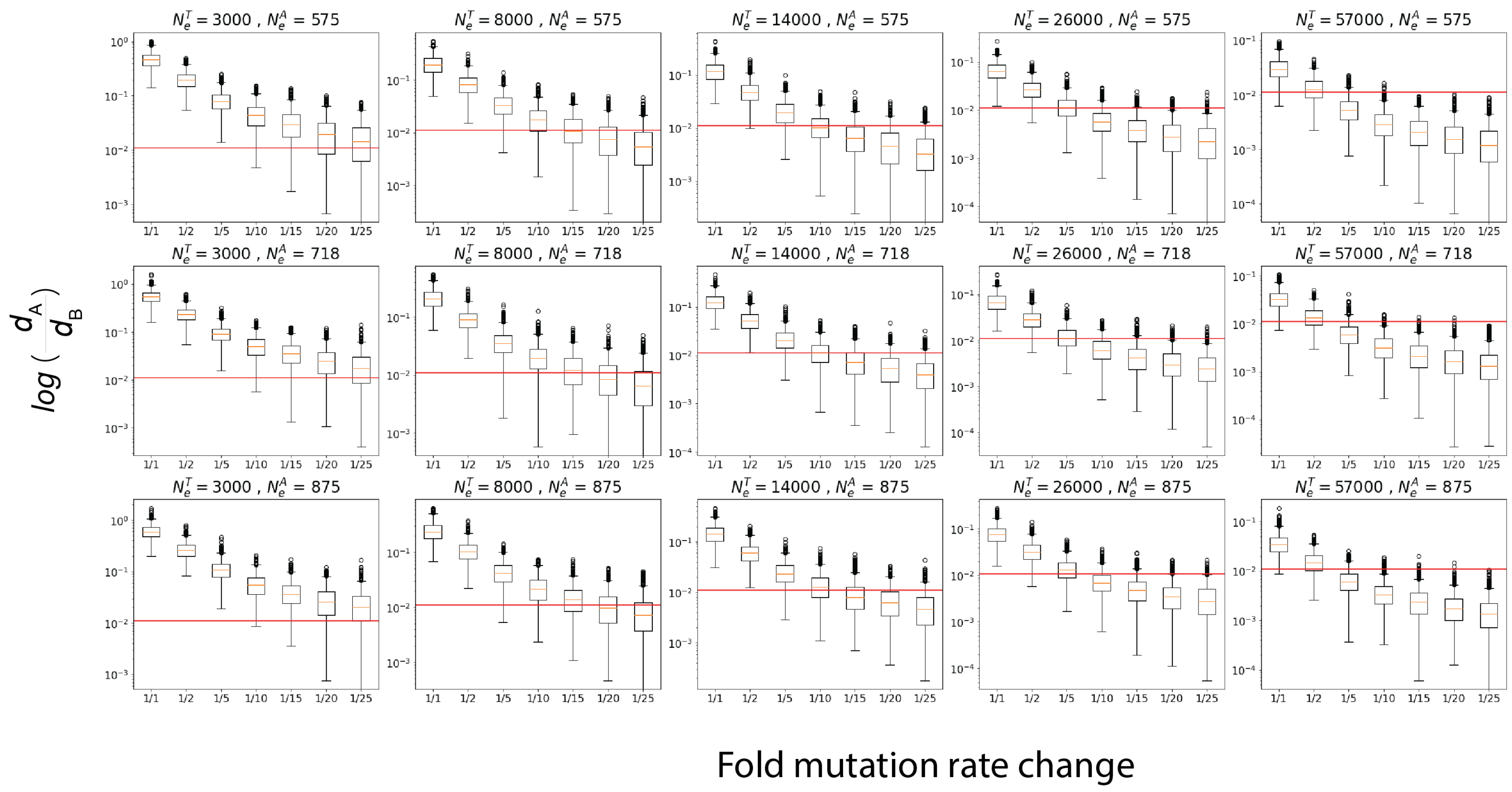


**Figure S7. Ancestry simulations over a range of population sizes and mutation rate changes (Related to Figure 3B,C).** Boxplot showing the degree of phylogenetic clustering in subpopulation A relative to subpopulation B over a range of mutation rate changes. Each set of simulations is repeated over a range of effective population sizes of subpopulation A ($N_e^A$) and a range of total effective population sizes ($N_e^T$) spanning the 95% confidence intervals estimated using BEAST assuming a constant population size tree prior. The panel in which $N_e^T$=8000 and $N_e^A$=718 recapitulates Figure 3B.

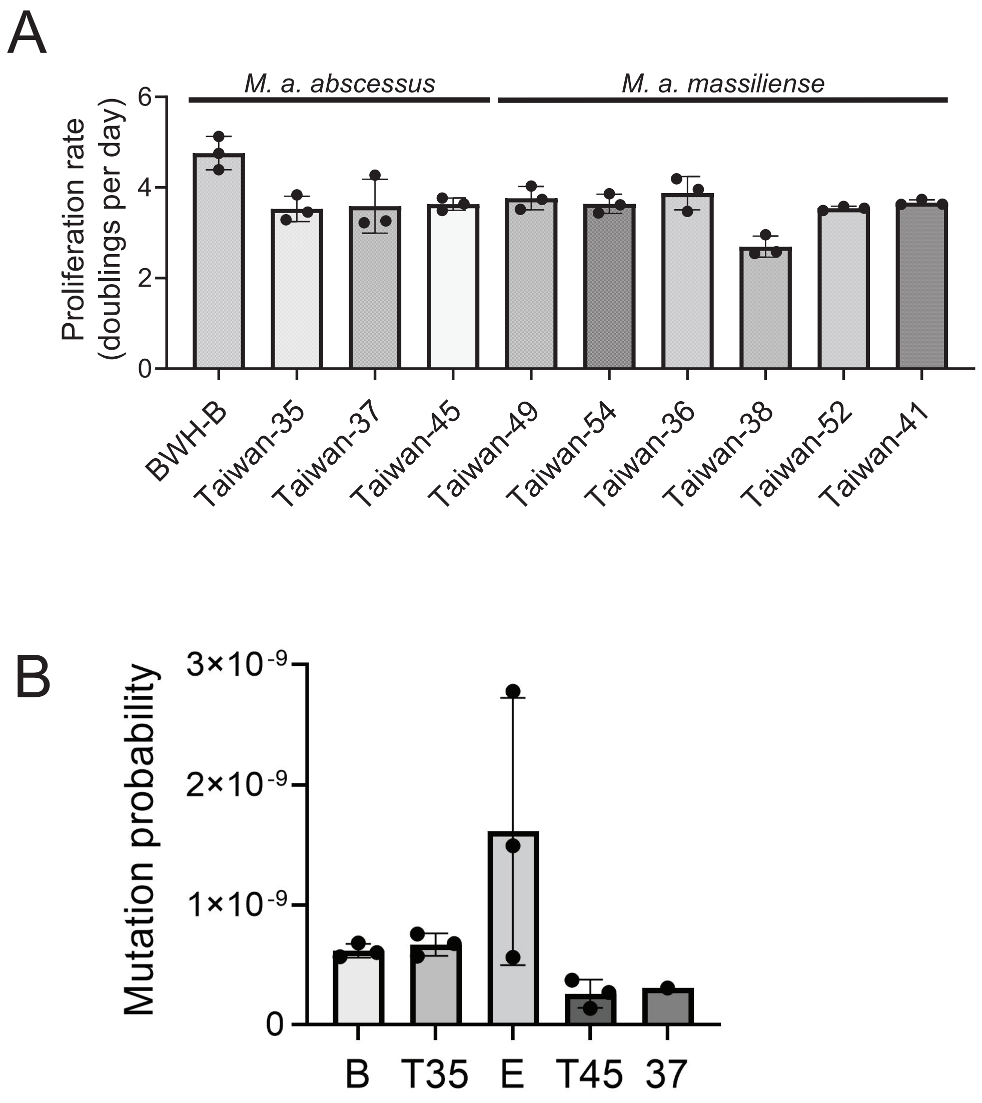


**Figure S8. Growth rates and mutation probability of *M. abscessus* strains. (Related to Figure 4C-E).** A) Growth rates of all isolates used in fluctuation assays. B) Probability of acquiring an amikacin resistance mutation per generation in *M. a. abscessus* strains including BWH-E. Data for BWH-B, Taiwan-35, Taiwan-37, and Taiwan-45 are reproduced from Figure 4C. Mean +/- SD for 3 biological replicates is displayed.

**Table S1**. Isolates used in this study

| Number of samples | BioProject ID | Citation |
|---|---|---|
| 854 | PRJEB2779 | Bryant, J et al. 2016. "Emergence and Spread of a Human-Transmissible Multidrug-Resistant Nontuberculous Mycobacterium." *Science* 354 (6313): 751–57. |
| 190 | PRJNA319839 | Hasan, NA et al. 2019. "Population Genomics of Nontuberculous Mycobacteria Recovered from United States Cystic Fibrosis Patients." *bioRxiv*. https://doi.org/10.1101/663559. |
| 173 | PRJEB7058 | Everall, I et al. 2017. "Genomic Epidemiology of a National Outbreak of Post-Surgical Mycobacterium Abscessus Wound Infections in Brazil." *Microbial Genomics* 3 (5): e000111. |
| 141 | PRJEB31559 | Doyle, RM et al. 2019. "Cross-Transmission Is Not the Source of New Mycobacterium Abscessus Infections in a Multi-Centre Cohort of Cystic Fibrosis Patients." *Clinical Infectious Diseases: An Official Publication of the Infectious Diseases Society of America*, June. https://doi.org/10.1093/cid/ciz526. |
| 69 | PRJNA420644 | Lipworth, S. *et al.* Whole-Genome Sequencing for Predicting Clarithromycin Resistance in Mycobacterium abscessus. *Antimicrob. Agents Chemother.* **63**, (2019) |
| 17 | PRJNA439313 | Yan, J. *et al.* Investigating transmission of Mycobacterium abscessus amongst children in an Australian cystic fibrosis centre. *J. Cyst. Fibros.* **19**, 219–224 (2020) |
| 11 | PRJNA297030 | Unpublished |
| 1 | PRJNA447908 | Unpublished |
| 1 | PRJNA495001 | Chhotaray, C. *et al.* Comparative Analysis of Whole-Genome and Methylome Profiles of a Smooth and a Rough Mycobacterium abscessus Clinical Strain. *G3* **10**, 13–22 (2020) |
| 1 | PRJEB1520 | Unpublished |
| 1 | PRJNA347845 | Fogelson, S. B. *et al.* Variation among human, veterinary and environmental Mycobacterium chelonae-abscessus complex isolates observed using core genome phylogenomic analysis, targeted gene comparison, and anti-microbial susceptibility patterns. *PLoS One* **14**, e0214274 (2019) |
| 1 | PRJNA566387 | Pearce, C. *et al.* Inhaled tigecycline is effective against Mycobacterium abscessus in vitro and in vivo. *J. Antimicrob. Chemother.* **75**, 1889–1894 (2020) |
| 1 | PRJNA401495 | Unpublished |

**Extended Data Table 2.** Description of Isolation Sources

| **Isolation Source** | **Number of samples (%)** |
|---|---|
| Pulmonary | 1181 (80.8%) |
| Skin and soft tissue swab or biopsy | 185 (12.7%) |
| Lymph node | 8 (0.5%) |
| Environmental | 4 (0.2%) |
| CSF | 3 (0.2%) |
| Blood | 2 (0.1%) |
| Feces | 2 (0.1%) |
| Clinical isolates from unknown source | 76 (5.2%) |

**Table S3.** Reference genomes used in this study

| **Strain** | **Citation** |
|---|---|
| *M. a. abscessus* ATCC 19977 | Ripoll, F et al. 2009. “Non Mycobacterial Virulence Genes in the Genome of the Emerging Pathogen Mycobacterium Abscessus.” *PloS One* 4 (6): e5660. |
| *M. a. massiliense* CCUG 48898=JCM 15300 | Sekizuka, T et al. 2014. “Complete Genome Sequence and Comparative Genomic Analysis of Mycobacterium Massiliense JCM 15300 in the Mycobacterium Abscessus Group Reveal a Conserved Genomic Island MmGI-1 Related to Putative Lipid Metabolism.” *PloS One* 9 (12): e114848. |
| *M. a. bolletii BD* | Yoshida M et al. 2018. “Complete Genome Sequence of a Type Strain of Mycobacterium Abscessus Subsp. Bolletii, a Member of the Mycobacterium Abscessus Complex.” *Genome Announcements* 6 (5). https://doi.org/10.1128/genomeA.01530-17. |

**Table S4.** Results from Bayesian Evaluation of Temporal Signal

| Model | Marginal likelihood (SD) |
|---|---|
| Relaxed clock, constant population size (with dates) | -7144301.64 (21.42) |
| Relaxed clock, constant population size (without dates) | -7144579.04 (21.01) |

Comparison of the marginal likelihoods with sampling dates and without sampling dates yields the following Bayes Factor (BF) calculation: $Log_{10}(BF) = 277.4$

**Table S5.** BEAST estimated Ne for the 'clustered' subpopulation A under a range of assumed clock rates

| Clock rate (SNPs/site/year) | $N_e^A$ [95% HPD] |
|---|---|
| $1.7x10^{-7}$ [a] | 718.99 [575.06, 875.16] |
| $8.5x10^{-7}$ | 142.79 [116.38, 175.52] |
| $1.7x10^{-6}$ | 71.75 [57.57, 87.07] |
| $8.5x10^{-6}$ | 14.27 [11.47, 17.35] |

[a] Mutation rate estimated for *M. a. abscessus* cluster 1 by Ruis et al.[13]

**Table S6.** Genotypes of UvrD/Rep family genes

| Gene name | Subspecies | Genome Position | Position in Gene | Ancestral Allele | Derived Allele |
|---|---|---|---|---|---|
| *MAB_1054* | *M. a. abscessus* | 1063899 | 2187 | A | G |
| *MAB_3515c* | *M. a. abscessus* | 3557910 | 2617 | G | C |
| *MAB_3515c* | *M. a. abscessus* | 3560378 | 149 | C | G |
| *MAB_3516c* | *M. a. abscessus* | 3562503 | 1197 | G | A |
| *MMASJCM_RS17495* | *M. a. massiliense* | 3533087 | 1737 | A | C |
| *MMASJCM_RS17515* | *M. a. massiliense* | 3538952 | 440 | A | G |

Gene names, genome positions, and position within genes are based on annoted reference genomes cited in Extended Data Table 2.

**Table S7.** Isolates used in fluctuation assays

| Isolate Name | Subspecies | Found in dominant cluster? | Genotype[a] | Isolation Source | Location |
|---|---|---|---|---|---|
| Mab-Taiwan-45 | *M. a. abscessus* | Yes | derived | Skin abscess | Taiwan |
| Mab-Taiwan-37 | *M. a. abscessus* | Yes | derived | Lung | Taiwan |
| Mab-BWH-E | *M. a. abscessus* | Yes | derived | Blood | United States |
| Mab-Taiwan-35 | *M. a. abscessus* | No | ancestral | Surgical wound site | Taiwan |
| Mab-BWH-B | *M. a. abscessus* | No | ancestral | Blood | United States |
| Mas-Taiwan-36 | *M. a. massiliense* | Yes | derived | Pleural effusion | Taiwan |
| Mas-Taiwan-38 | *M. a. massiliense* | Yes | derived | Ascites | Taiwan |
| Mas-Taiwan-41 | *M. a. massiliense* | Yes | derived | Breast abscess | Taiwan |
| Mas-Taiwan-52 | *M. a. massiliense* | Yes | derived | Ear (otorrhea) | Taiwan |
| Mas-Taiwan-49 | *M. a. massiliense* | No | ancestral | Eye | Taiwan |
| Mas-Taiwan-54 | *M. a. massiliense* | No | ancestral | Ear (external) | Taiwan |

[a] Full genotypes are listed in Extended Data Table 4.

**Table S8.** Baseline Rates of Resistance to Amikacin in Clinical Isolates

| Strain | Fraction of cells resistant to amikacin |
|---|---|
| Mas-Taiwan-36 | n.d. |
| Mas-Taiwan-38 | $8.77 \times 10^{-8}$ |
| Mas-Taiwan-41 | n.d. |
| Mas-Taiwan-52 | $1.59 \times 10^{-8}$ |
| Mas-Taiwan-49 | n.d. |
| Mas-Taiwan-54 | $5 \times 10^{-8}$ |
| Mab-BWH-B | $1 \times 10^{-8}$ |
| Mab-BWH-E | $1 \times 10^{-8}$ |
| Mab-Taiwan-35 | $1 \times 10^{-8}$ |
| Mab-Taiwan-37 | n.d. |
| Mab-Taiwan-45 | n.d. |

n.d. = no resistant colonies were detected

**List of Supplementary Data Files:**

1. mabsc_all_sample_metadata.csv: Accession IDs and metadata for all isolates used in this study.
2. MAB_homoplasies_enriched_dominant_clusters_nonsynonymousOnly.csv: Full list of nonsynonymous mutations ranked by their enrichment in *M. a. abscessus* DCs
3. MAB_homoplasies_enriched_dominant_clusters.csv: Full list of all mutations (synonymous or nonsynonymous) ranked by their enrichment in *M. a. abscessus* DCs
4. MAB_homoplasies_enriched_all_clusters_nonsynonymousOnly.csv: Full list of all nonsynonymous mutations ranked by their enrichment in all *M. a. abscessus* clusters
5. MAB_homoplasies_enriched_all_clusters.csv:Full list of all mutations (synonymous or nonsynonymous) ranked by their enrichment in all *M. a. abscessus* clusters
